# Characterization of atypical BAR domain-containing proteins coded by *Toxoplasma gondii*

**DOI:** 10.1101/2024.06.13.598837

**Authors:** Noha Al-Qatabi, Maud Magdeleine, Sophie Pagnotta, Amélie Leforestier, Jéril Degrouard, Ana Andreea Arteni, Sandra Lacas-Gervais, Romain Gautier, Guillaume Drin

## Abstract

*Toxoplasma gondii*, the causative agent of toxoplasmosis, infects cells and replicates inside *via* the secretion of factors stored in specialized organelles (rhoptries, micronemes, dense granules) and the capture of host materials. The genesis of the secretory organelles and the processes of secretion and endocytosis depend on vesicular trafficking events whose molecular bases remain poorly known. Notably, there is no characterization of the BAR (Bin/Amphiphysin/Rvs) domain-containing proteins expressed by *T. gondii* and other apicomplexans, although such proteins are known to play critical roles in vesicular trafficking in other eukaryotes. Here, by combining structural analyses with *in vitro* assays and cellular observations, we have characterized *Tg*REMIND (REgulators of Membrane Interacting Domains), involved in the genesis of rhoptries and dense granules, and *Tg*BAR2 found at the parasite cortex. We establish that *Tg*REMIND comprises an F-BAR domain that can bind curved neutral membranes with no strict phosphoinositide requirement and exert a membrane remodeling activity. Next, we establish that *Tg*REMIND contains a new structural domain called REMIND, which negatively regulates the membrane-binding capacities of the F-BAR domain. In parallel, we report that *Tg*BAR2 contains a BAR domain with an extremely basic membrane-binding interface able to deform anionic membranes into very narrow tubules. Our data show that *T. gondii* codes for two atypical BAR domain-containing proteins with very contrasting membrane-binding properties, allowing them to function in two distinct regions of the parasite trafficking system.

## Introduction

Apicomplexan parasites are highly polarized cells with a secretory system that includes unique apical organelles called the micronemes and rhoptries, but also dense granules, which are more dispersed throughout the cytosol. All these organelles contain protein factors that are essential for parasite virulence. Micronemal proteins (MIC) guarantee parasite motility and host cell attachment and invasion, whereas rhoptry (ROP) and dense granule (GRA) proteins play a pivotal role in the creation and maintenance of the parasitophorous vacuole (PV), in which the parasite develops inside a host cell, and in altering the immune response and metabolic state of the infected cell. Given the critical roles of these factors, elucidating the molecular mechanisms by which these secretory organelles are formed is of prime interest. For this purpose, *T. gondii* is a powerful model for studying protein trafficking and organelle biogenesis in apicomplexan parasites.

The synthesis, sorting, and transport of MIC, ROP, and GRA proteins are tightly linked to the biogenesis of the secretory organelles. In *T.gondii*, the genesis of micronemes and rhoptries starts with the budding of vesicles containing ROP and MIC proteins synthesized in the form of propeptides from the trans side of the Golgi apparatus. These vesicles fuse with an endosomal compartment that serves as an intermediate compartment for proteins destined for secretion and not, as in other eukaryotic cells, for recycling or degradation of endocytosed material. It is assumed that MIC proteins transit from an early to a late compartment of this endosomal-like compartment (ELC) to undergo processing by specialized proteases originating from the parasite vacuole, to be transferred to the micronemes in an active form (1). ROP proteins are assumed to transit directly from the early endosomal compartment to pre-rhoptries to be processed into functional factors. Pre-rhoptries, which are large vesicles situated between the Golgi and the apical region of the parasite, undergo a process of condensation and elongation before cytokinesis, eventually developing into mature rhoptries (1). In contrast, dense granules are directly formed from the TGN in a clathrin-dependent manner and exhibit characteristics of both the constitutive and regulated secretory pathways observed in mammalian cells (2).

Much evidence suggests that the trafficking system of Apicomplexan is divergent from those found in other eukaryotic cells like human or yeast cells, but it also relies on a more limited number of components. For instance, *T. gondii* only depends on a basic set of small G-proteins Rabs (3) and lacks highly conserved endocytic factors like the components of Endosomal Sorting Complexes Required for Transport (ESCRT) or Golgi-associated, Gamma adaptin ear containing, Arf binding coat proteins (GGAs) that regulate the anterograde protein transport between the *trans*-Golgi network and the endo-lysosomal compartments (4). Nevertheless, many standard components of the vesicle budding, transport and fusion machinery have been identified, including N-ethylmaleimide-sensitive fusion (NSF) factor (5), Soluble N-ethylmaleimide-sensitive-factor Attachment protein Receptor (SNAREs) proteins (6), subunits of the coatomer (7), clathrin adaptor complexes (8), Vacuolar Protein Sorting (VPS) proteins (9,10) and dynamin-related protein B (11). In addition, evolutionarily-conserved tyrosine and leucine-based motifs functioning in the transport and sorting of MIC and ROP protein have been characterized (12,13). What is possibly the most fascinating is that many trafficking factors known to have given roles in well-described cellular systems are repurposed in *T. gondii* for different functions. Typically, *Tg*SORTLR, clathrin-adaptor protein 1 *Tg*AP1, and dynamin-related protein *Tg*DrpB function together at the Golgi to sort MIC and ROP protein. In contrast, their human counterparts exert their function in the endocytotic/endosomal system.

Bin/Amphiphysin/Rvs (BAR) domain-containing proteins play pivotal roles in different stages of vesicular trafficking in eukaryotic cells, particularly endocytosis, but can also act as membrane remodelers (14). BAR domains form crescent-shaped homodimers that can bind and often bend membranes and, in some cases, favor membrane scission. Depending on their shape, size, and combination with membrane-binding amphipathic helices (AH), these BAR domains are classified as BAR, N-BAR, F-BAR, or I-BAR. They impart many proteins with the capacity to sense and/or change membrane geometries. It is important to note that the behavior of these BAR-domain-containing proteins intimately depends on the features of the membrane to which they bind, such as their shape but also negative charge density and tension; in some cases, the membrane-binding capacity of the BAR-containing proteins can be self-regulated or controlled by an external partner. Most BAR-domain-containing proteins integrate other domains or motifs, enabling them to play many functions in different subcellular regions.

At least two *T. gondii* proteins have been listed as encompassing a BAR domain (15). The first protein (TGME49_259720), predicted to have an N-terminal F-BAR domain, has been found recently to localize to internal compartments in *T.gondii*, including rhoptries, dense granules, Golgi, and post-Golgi compartments (16,17). Remarkably, the genetic ablation of this protein, termed *Tg*REMIND, results in the absence of dense granules and the formation of abnormal transparent rhoptries, leading to severe inhibition of the parasite’s motility, host invasion and dissemination capabilities (17). *Tg*REMIND is also predicted to contain a C-terminal domain termed REMIND with a totally new fold and unknown function. Moreover, it has been proposed that *Tg*REMIND recognizes phosphoinositides (PIPs) *via* its F-BAR and REMIND domains to target membranes. Finally, the protein has been reported to co-immunoprecipitate with plenty of trafficking factors (17). Collectively, these data suggest that *Tg*REMIND might have an essential role in membrane trafficking and the formation of secretory organelles required for infection by *T.gondii*.

The second putative BAR domain-containing protein (TGME49_320760, called *Tg*BAR2) is localized at the cell periphery, notably at the IMC (16). It is still very unclear how *T. gondii* endocytoses materials. A circular invagination of the PM with a dense neck, termed the micropore, is believed to be the essential structure related to this process. Recent studies have revealed the existence of a network of proteins with homology with endocytic proteins responsible for the scaffolding of micropores but did not identify *Tg*BAR2 as a component of this network (16,18). Except for these few data, we have no insights into the structure and the functional traits of these two potential BAR domain-containing proteins.

To get better insights into the potential function of these two proteins, we have analyzed their structural features and membrane-binding properties *in vitro* and in a cellular context. We establish that *Tg*REMIND has an F-BAR domain that can preferentially associate with curved neutral membranes with no strict PIPs requirement. It has the intrinsic capacity to deform and potentially disrupt the membranes. Next, we show that the REMIND domain constitutes a novel kind of structural domain that tunes the ability of *Tg*REMIND to associate with the membrane *via* its F-BAR domain. These data explain the ubiquitous localization of *Tg*REMIND within the parasite and suggest it is potentially a membrane-remodeling device under the control of the REMIND domain. In parallel, we report that *Tg*BAR2 contains a BAR domain that distinguishes itself from well-known BAR domains by its extremely basic membrane-binding interface. The full-length protein targets preferentially anionic membranes, which might explain its localization at the cell periphery. It can strongly remodel the membranes and even form micellar tubules. Collectively, our data indicate that *T. gondii* codes for two atypical BAR domain-containing proteins with very contrasting features in terms of membrane targeting and remodeling power.

## Results

### *Tg*REMIND has an F-BAR domain to bind neutral membranes with no strict PIP requirement

The 80-348 region of *Tg*REMIND is predicted to correspond to a Fes/CIP4 homology-Bin/Amphiphysin/Rvs (F-BAR) domain **(Fig. 1A**, (17)). **Figure 1B** shows a tridimensional model of this region, established by AlphaFold, which has the features of a canonical BAR: two monomers with three long α-helices arranged into a twisted coiled-coil define a six-helix bundle dimer with a crescent shape. The domain’s concave face displays small patches of positive and negative charges instead of being highly basic, as found using electrostatic potential mapping (**Fig. 1C**). We expressed the 70-347 region of *Tg*REMIND appended with an N-terminal His-tag in *E. coli*; this construct was then first isolated by affinity chromatography, and then further purified by size-exclusion chromatography (SEC). Endogenous cysteine residues, predicted to be exposed at the surface of the F-BAR domain, were substituted by alanine or serine residues to limit the aggregation of this construct during purification (**Sup.** Fig. 1A). Gel-filtration analyses suggested that the purified protein was mainly in a trimer of dimers form or in a monomeric form (∼190 kDa or ∼37 kDa, respectively). By circular dichroism (CD) spectroscopy, we established that it was folded correctly with a percentage of α-helix (74%) in excellent agreement with the percentage estimated from the AlphaFold model **(Fig. 1D)**. All these data strongly suggest that the N-terminal region of *Tg*REMIND contains an F-BAR domain.

**Figure 1.**
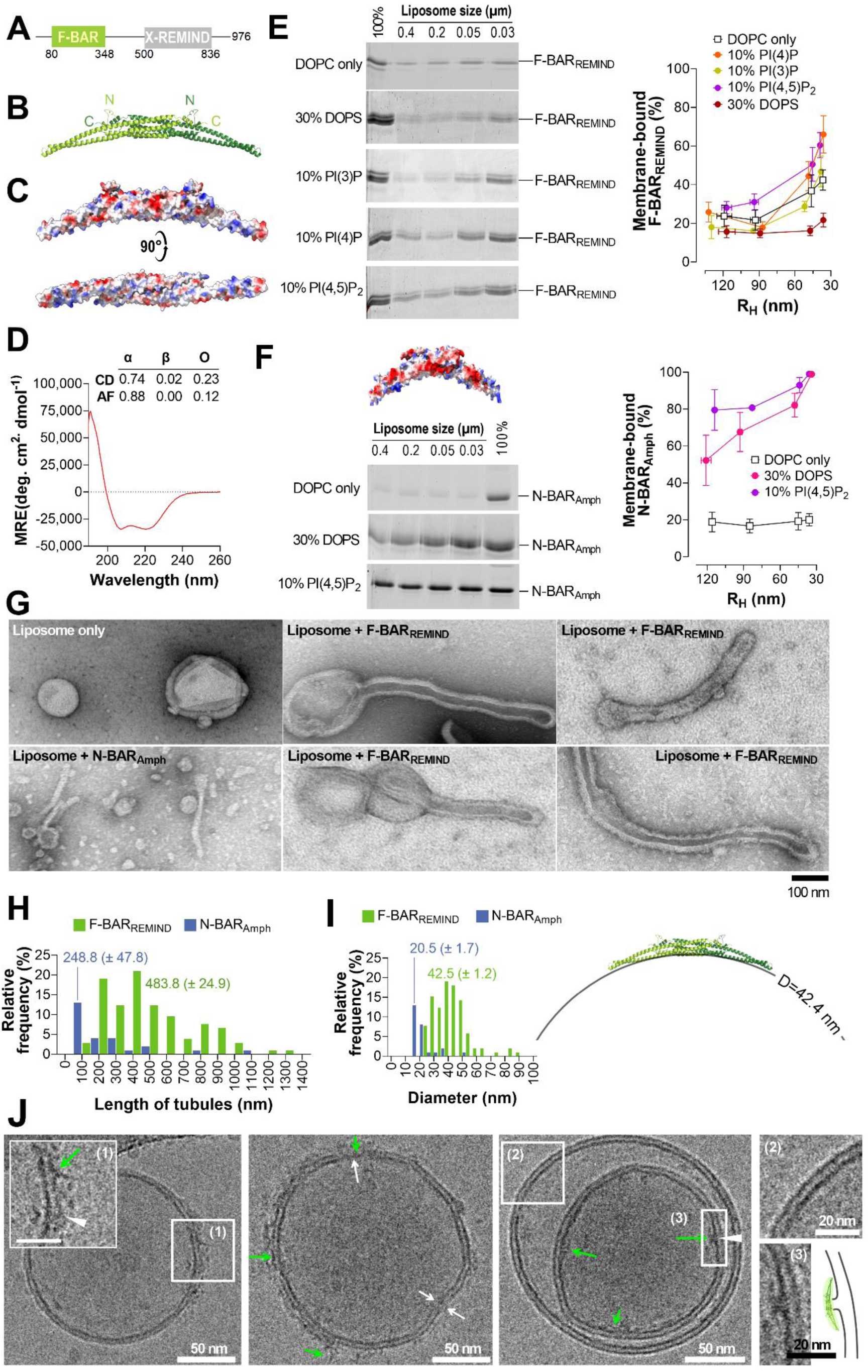
*Tg*REMIND F-BAR domain binds membrane with no strict PIP requirement. **(A)** Predicted structural organization of *Tg*REMIND. **(B**) Ribbon representation of the AlphaFold-predicted model of the F-BAR domain of *Tg*REMIND in a dimeric form. The position of each monomer’s N- and C-terminal ends is indicated. **(C)** Same view as (B) or view of the concave face of the F-BAR domain with the surface colored by electrostatic potential (red = -16.9 kTe^-1^, blue = +16.9 kTe^-1^). **(D)** Far-UV CD spectrum of purified F-BAR_REMIND_ (3 µM) in 20 mM Tris, pH 7.4, 120 mM NaF buffer recorded at room temperature. The percentage of α-helix, β-sheet, and other structures, derived from the CD spectra analysis and the AlphaFold-predicted model obtained using the DSSP algorithm, is given. MRE: mean residue ellipticity, H: α-helix, E: β-sheet, O: other structures. (**E**) Flotation assay. F-BAR_REMIND_ (0.75 µM) was incubated with liposomes (750 µM total lipids) only made of DOPC and additionally containing 30% DOPS or 10% PIPs (PI(3)P, PI(4)P or PI(4,5)P_2_ at the expense of DOPC), extruded through pores of defined size, indicated at the top of the gel. This incubation was performed for 1 h at 25 °C in 50 mM Tris, pH 7.4, 150 mM NaCl (TN) buffer under constant agitation. After centrifugation, the liposomes were recovered at the top of sucrose cushions and analyzed by SDS-PAGE. The amount of membrane-bound F-BAR was determined using the content of the first lane (100% total) as a reference based on the SYPRO Orange signal. The percentage of membrane-bound F-BAR_REMIND_ is represented as the function of the average hydrodynamic radius (R_H_) of liposomes. Data are expressed as mean ± s.e.m. (n = 3-5). **(F)** Flotation assay. N-BAR domain of human amphiphysin (N-BAR_Amph_, 0.75 µM) was incubated with liposomes (750 µM lipids), only made of DOPC and additionally containing 30% DOPS or 10% PI(4,5)P_2_, extruded through pores of defined size, for 1 h at 25 °C in 50 mM Tris, pH 7.4, 150 mM NaCl (TN) buffer. The percentage of membrane-bound N-BAR_Amph_ is shown as the function of the radius of liposomes (n = 3). (**G**) Negative-staining EM. Liposomes made of Folch fraction I lipids (30 µM lipids) and extruded through 0.4 µm pores were incubated with F-BAR_REMIND_ or N-BAR_Amph_ (1.9 µM) for 2 h at room temperature. A control picture of liposomes alone is shown. Scale bar = 100 nm. (**H**) Length distributions of membrane tubules induced by F-BAR_REMIND_ and N-BAR_Amph_ (n =105 and 26, respectively). **(I)** Diameter distribution of tubules and N-BAR_Amph_ (n =105 and 26, respectively). The structure of the F-BAR domain of *Tg*REMIND seems adapted to the diameter of tubules that have been experimentally measured. **(J)** Cryo-EM. Folch fraction liposomes (90 µM lipids), extruded through 0.4 µm pores, were mixed with F-BAR_REMIND_ (6 µM) at P/L= 1/15 and dialyzed three times under constant stirring in TN buffer for 30 min at cold temperature. Enlargements: (1) Outer leaflet destabilization (white arrowhead) by F-BAR_REMIND_, (2) Intact bilayer, (3) Outer leaflet destabilization (white arrowhead) by F-BAR_REMIND_. Green arrows point to some individual F-BAR_REMIND_ molecules. White arrows indicate local membrane disruption (both leaflets)

Next, we tested the capacity of this construct, referred to as F-BAR_REMIND,_ to bind membranes with given curvature and lipid composition using flotation assays. First, we mixed F-BAR_REMIND_ with 1,2-dioleoyl-*sn*-glycero-3-phosphocholine (DOPC) liposomes of different sizes (from ∼130 to 35 nm in radius) obtained by sequential extrusion. Liposomes were isolated at the top of sucrose cushions after centrifugation, and membrane-bound protein was quantified using SDS-PAGE (19,20). F-BAR_REMIND_ was slightly associated (23.7% total protein) with large liposomes (with a hydrodynamic radius, R_H_ ∼ 120 nm) and more with smaller ones (up to 42.4% with the smallest liposomes, R_H_ < 40 nm, **Fig. 1E**). This suggested that it could bind neutrally-charged membranes in a curvature-dependent manner. Next, we tested liposomes of different sizes containing 30% of 1,2-dioleoyl-*sn*-glycero-3-phospho-L-serine (DOPS) at the expense of DOPC. We observed that F-BAR_REMIND_ had a much lower avidity for these liposomes but still showed a curvature-dependent association. The fraction of membrane-bound protein increased from 15.6 to 21.7% as the liposome size decreased from 117 to 36 nm. Furthermore, we tested F-BAR_REMIND_ with liposomes that contained 10% of either PI(3)P, PI(4)P, or PI(4,5)P_2_. We found it associated more strongly with small liposomes (36 nm) enriched with PI(4)P or PI(4,5)P_2_ (up to 66% of total protein) than with small liposomes only composed of DOPC.

In parrallel, we tested an archetypical BAR domain, *i.e.*, the N-BAR domain of human amphiphysin ((21,22), residues 2–242, N-BAR_Amph_) with liposomes made of DOPC only or enriched with DOPS. We observed that it bound weakly to DOPC membranes, whatever their curvature, but strongly associated with DOPS-rich membranes in a slightly curvature-dependent manner (**Fig. 1F**), in line with results obtained by Peter *et al.* (21). These results suggest that F-BAR_REMIND_ does not interact with curved membranes through electrostatic interactions like N-BAR_Amph_ and many other BAR domains. Instead, it can substantially bind curved, neutrally charged membranes, as observed for the F-BAR domains of the yeast proteins Rdg1p, Bzz1p, and Hop1p (**Fig.S1C**, (23)), with an avidity that can be slightly enhanced by PIPs, as also observed for Rdg1p.

### F-BAR_REMIND_ domain can remodel and destabilize membranes

Having determined that F-BAR_REMIND_ is adapted to curved membranes, we investigated whether it could deform flat membranes as observed with other BAR-domain-containing proteins by negative staining electron microscopy (EM) (24–28). We incubated this construct with large liposomes composed of Folch fraction I lipids (containing PS and PIPs) at different protein-to-lipid (P/L) molar ratios, ranging from 1/120 to 1/15. Comparatively to observations carried out with liposomes alone, we observed tubules emanating from many liposomes when incubated with F-BAR_REMIND_ (**Fig. 1G)** with one, sometimes two, tubules *per* liposome **(Fig.S1D)**. Tubulation was observed preferentially at high P/L ratios (1/15), suggesting that this process was dependent on the density of protein on the membrane surface **(Fig. S1E)**. The length of tubules, estimated from a large set of EM pictures obtained through replicated assays with different F-BAR_REMIND_ batches, was highly variable, with an average value of 483.8 ± 50 nm (**Fig. 1H**). Comparatively, the diameter of tubules was very homogeneous, with an average value of 42.5 ± 1.4 nm (**Fig. 1I**). This was remarkably consistent with our estimate that the concave face of F-BAR_REMIND_ would fit a membrane tubule of 42.4 nm in diameter. Comparative tests showed that N-BAR_Amph_ induced the formation of narrower tubules **(Fig. 1G-I)**. Overall, according to our structural predictions, our results indicate that *Tg*REMIND contains a *bona fide* F-BAR domain able to induce the formation of tubules with a diameter similar to that observed for tubules induced by other F-BAR-domain-containing proteins (27,29).

To get confirmation of this phenomenon and a higher-resolution view of the membrane-binding and remodeling capacity of F-BAR_REMIND_, we examined liposomes in the presence or absence of F-BAR_REMIND_ (at P/L= 1/15) by cryo-EM. We observed that the F-BAR_REMIND_ domain decorated the surface of liposomes and modified the bilayer but did not form tubules in our experimental conditions (**Fig.1J**, green arrows). Interestingly, we observed that the F-BAR_REMIND_ interacted with the outer or inner leaflet of the membrane and, in some areas, disrupted either or both leaflets (**Fig.1J**, white arrowheads, and arrows, respectively). Moreover, we also observed some proteins within the interior of the liposomes, confirming that the integrity of the membrane was lost (**Fig.1J**, white arrow). Collectively, these data show the capacity of the F-BAR_REMIND_ to interact with membranes, locally modify their shape, and potentially disrupt them.

### Impact of charge reversal mutations on the membrane-binding properties of F-BAR_REMIND_

We mentioned above the resemblance between F-BAR_REMIND_ and the F-BAR domain of Rgd1p regarding membrane-binding capacities. Interestingly, the Rdg1p F-BAR domain can make specific contacts with the polar head of PIPs *via* a cluster of basic residues localized in its concave face (23). We identified an analogous, albeit smaller, cluster in F-BAR_REMIND_ composed of the K210, R214, R217, and R250 residues **(Fig. S2A, Fig. 2A)**. Therefore, we designed three mutants in which two out of these four residues were substituted by anionic residues (R214E/R217E, R217E/R250E, K210D/R214E). This markedly impacts the electrostatic features of the concave face of the F-BAR domain (**Fig. 2B**). We purified the corresponding mutants **(Fig. S2A, B)** and assessed that they were adequately folded by CD spectroscopy **(Fig. S2C)**. Using flotation assays, we found that, overall, these mutated forms of F-BAR_REMIND_ bound only slightly less to PI(4,5)P_2_-containing liposomes compared to the wild-type form, notably when these liposomes were large **(Fig.2C)**. The most defective mutant was the R214E/R217E mutant, suggesting that R214 and R217 residues could be primarily involved in recognizing PIPs. Because F-BAR_REMIND_ substantially binds to neutral membranes, we also examined the potential role of hydrophobic interaction in the membrane-binding process. We identified a portion of the lateral side of F-BAR as highly hydrophobic and substituted two hydrophobic residues with anionic ones to obtain an L201D/M212E mutant. Albeit well-folded (**Fig. S2A-E)**, this construct was impaired in its capacity to bind liposomes, whatever their size, comparatively to the wild-type form of F-BAR_REMIND_ **(Fig.2D)**. However, EM observation showed that all these mutants deformed liposomes in tubulation assays at a high P/L ratio **(Fig. 2E)**. We conclude that the interaction of F-BAR_REMIND_ with the membrane is not primarily dictated by specific interaction with PIPs but most probably by a combination of hydrophobic and short-range electrostatic interaction.

**Figure 2.**
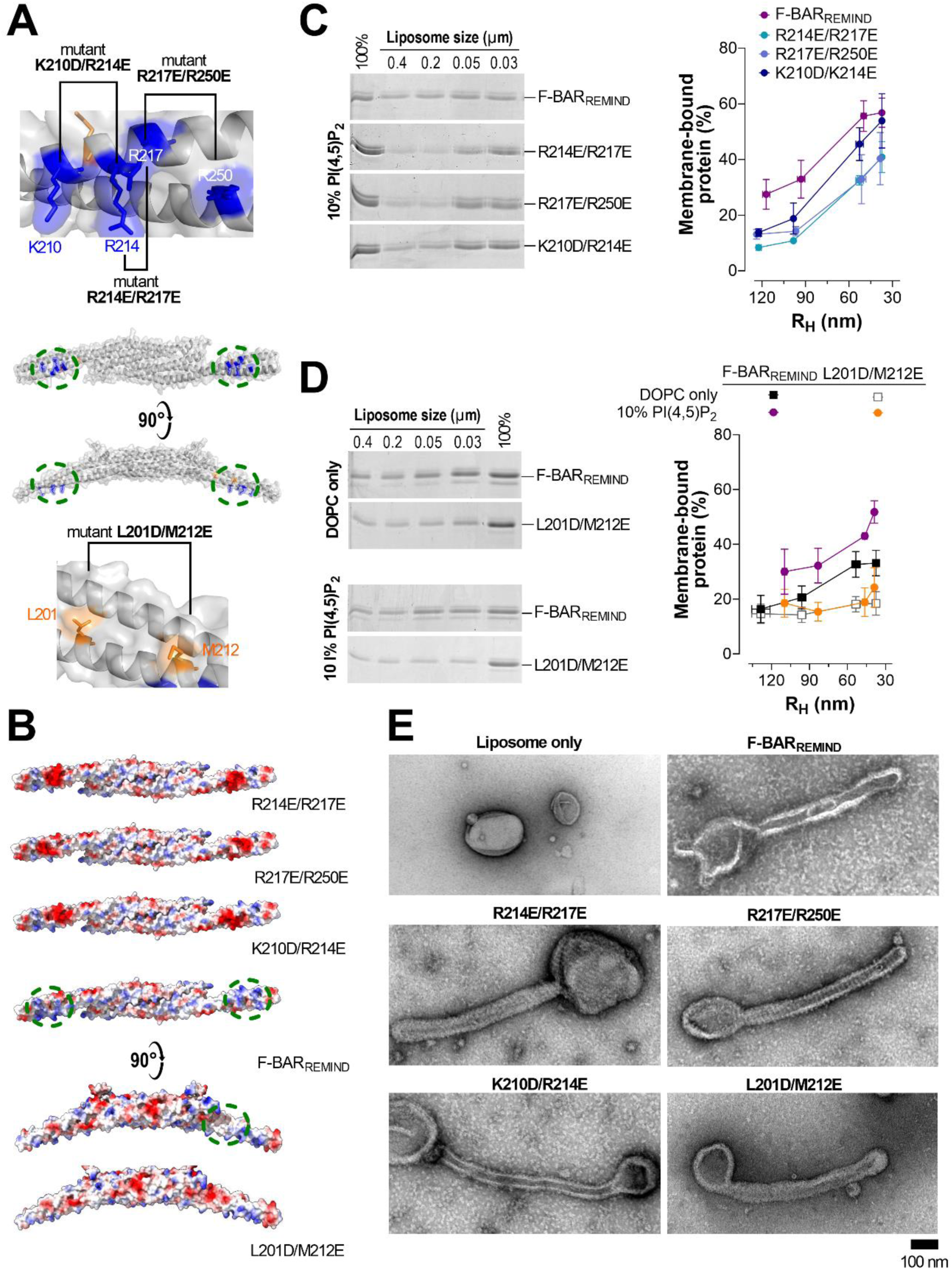
Charge reversion in the concave and lateral side of F-BAR_REMIND_ affects how it binds to and deforms membranes. **(A)** Localisation of arginine and lysine residues that constitute a basic cluster in the concave face of F-BAR_REMIND_ (in blue) and localization of solvent-exposed hydrophobic residues in the lateral side of this domain (in orange). These residues were substituted by anionic residues (aspartate or glutamate) to test their contribution to the F-BAR_REMIND_/membrane interaction. **(B)** Electrostatic features of the molecular surface of F-BAR_REMIND_ and its mutated forms (red = -16.9 kTe^-1^, blue = +16.9 kTe^-1^). **(C)** Flotation assay. F-BAR_REMIND_ or K210D/R214E, R214E/R217E or R217E/R250E mutant (0.75 µM) was incubated with liposomes composed of DOPC/PI(4,5)P_2_ (90:10, 750 µM lipids), with a defined radius, for 1 h at 25 °C. Data are represented as mean ± s.e.m. (n = 3-5). **(D)** Flotation assay. F-BAR_REMIND_ or L201D/M212E mutant (0.75 µM) was incubated with liposomes (750 µM) composed of DOPC or DOPC/PI(4,5)P_2_ (90:10), with a defined radius, for 1 h at 25 °C. Mean ± s.e.m. (n = 3-4). **(E)** Negative-staining EM. Liposomes made of Folch fraction I lipids (30 µM) were incubated with F-BAR_REMIND_ or its mutated form (1.9 µM) for 2 h at room temperature (scale bar = 100 nm). A control picture of liposomes only is shown

### REMIND domain is a new structural domain without any membrane-binding capacity

The 500-840 region of the *Tg*REMIND protein is predicted by AlphaFold to be composed of fifteen α-helices and three β-sheets (**Fig. 3A**). This domain, referred to as REMIND, has no homology with any other known structural domains (17). We examined whether this predicted structure remained stable in water during a 1-µs long molecular dynamics (MD) simulation. As shown in **Fig. 3B**, we observed no unfolding of the structure over time, with a root-mean-square deviation (RMSD) inferior to 0.7 nm. By calculating the root-mean-square fluctuation (RMSF) for each Cα of the protein backbone (**Fig. 3C**), we identified that the N-and C-terminal ends of the protein were mobile, as were three intrinsically disordered linkers between α1 and α2 helices, α2 and α3 helices, and α8 helix and β3 sheet. We also noted that a couple of α-helices (α10 and α11) were mobile yet without getting unfolded. Collectively, these data suggest that the region 500-840 of *Tg*REMIND corresponds to a stable structural domain.

**Figure 3.**
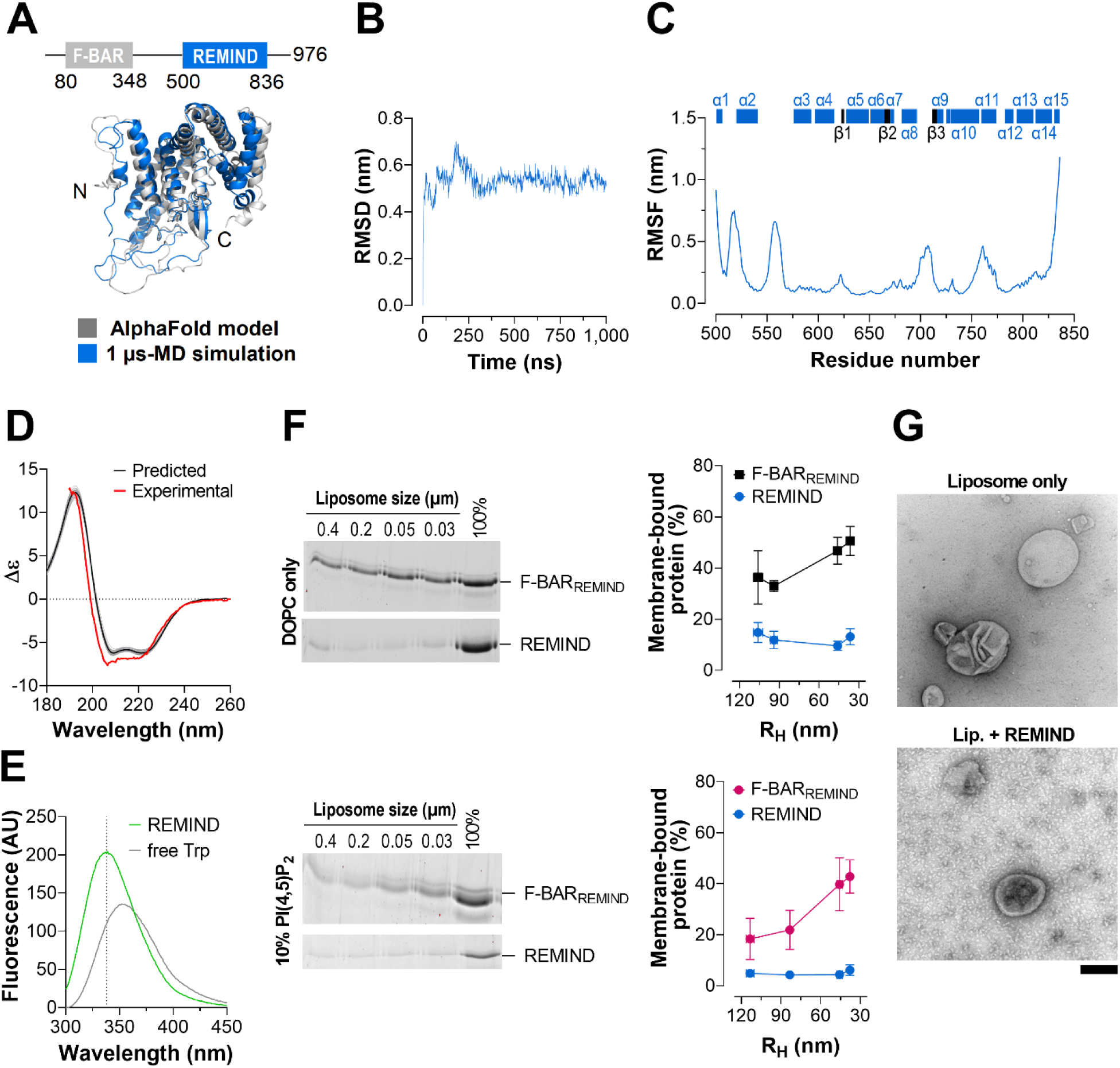
REMIND region encompasses a folded domain with no membrane-binding capacity. (**A**) Ribbon representation of the three-dimensional model of the REMIND domain (region 500-836 of *Tg*REMIND) established by AlphaFold (in grey) and of the final conformation of this domain after 1 µs-MD simulation in water (in blue). The N- and C-terminal ends of the domain are indicated. (**B**) RMSD of the Cα atoms with respect to the starting and equilibrated structure of the REMIND domain as a function of time. **(C)** RMSF values of atomic positions of Cα atoms, indicative of the internal protein motions, are shown as a function of residue number. The localization of α-helix and β-sheet along the sequence is indicated. **(D)** Predicted and experimental CD spectra of the REMIND domain. Spectra were predicted from configurations of the REMIND domain collected every 10 ns during the MD simulation (grey spectra) using the PDBMD2CD algorithm. An average spectrum is represented in black. For comparison, the far-UV CD spectrum of purified *Tg*REMIND[495-840] construct (REMIND, 2 µM) in 20 mM Tris, pH 7.4, 120 mM NaF buffer at room temperature is shown (in red). **(E)** The intrinsic fluorescence of REMIND (1µM) was measured in TN buffer at 30 °C. A spectrum was measured with free L-tryptophane (4 µM) as a comparison **(F)** Flotation assay. REMIND (0.75 µM) was incubated with liposomes (750 µM lipids) of different mean radii, composed of DOPC or DOPC/PI(4,5)P_2_ (90:10) for 1 h at 25 °C under agitation. Data are represented as mean ± s.e.m (n = 3-4). **(G)** Negative staining-EM images of liposomes (30 µM lipids), composed of Folch fraction I lipids, mixed or not with REMIND (1.9 µM). Scale bar = 500 nm.

To further demonstrate this, we purified the [495-840] region of *Tg*REMIND comprising the REMIND domain to experimentally determine its structure (**Fig.S3)**. Unfortunately, attempts to crystallize this protein were unsuccessful. We therefore performed CD measurements to determine whether this construct was folded and, if true, to estimate its secondary structure content. We obtained a spectrum with two minima at 206-208 nm and 222 nm, indicating that the construct was well-folded (**Fig. 3D**). Then, we compared this spectrum with spectra generated from a series of 50 conformations adopted by the predicted REMIND domain during the one µs-MD simulation and found an excellent match. The CD spectrum analysis indicated that the construct is mainly helical (up to 60% helix) with a low β-sheet content in agreement with the corresponding values inferred from the tri-dimensional model. Additionally, we measured the intrinsic fluorescence of the REMIND domain by recording the emission fluorescence spectrum of tryptophan residues (eight in the purified construct). A maximum fluorescence intensity was measured at λ = 338 nm **(Fig. 3E)**. In comparison, we obtained a maximum fluorescence intensity at λ = 352 nm with free L-tryptophane in solution. These data indicated that some tryptophane residues of REMIND are buried in a hydrophobic environment, as predicted by the AlphaFold model. Jointly, our data establish the REMIND domain as being a new kind of structural domain whose folding probably corresponds to that predicted *in silico*.

Next, we tested whether the REMIND domain could bind to membranes. This construct (0.75 µM) was mixed with liposomes (750 µM lipids) of various sizes, either only composed of DOPC or doped with 10% PI(4,5)P_2_. We found that REMIND did not bind to these liposomes, whatever their composition and curvature, in stark contrast to F-BAR_REMIND_, used as a positive control (**Fig. 3F, G)**; moreover, we observed that REMIND, even at high P/L (1/30), could not deform liposomes. At odds with previous results based on protein-lipid overlay assays (17), our data suggest that the REMIND domain has no ability to bind membranes and instead has another function.

### F-BAR_REMIND_ but not REMIND domain can bind endomembranes

To strengthen our conclusions based on *in vitro* assays while also getting more insights into the features of F-BAR_REMIND_ and REMIND domains, we analyzed whether over-expressing these domains, separately or together in full-length *Tg*REMIND, induced membrane remodeling in a cellular context. Such a strategy was used to define better the functional traits of diverse BAR domain proteins (21,30–32). Because the transgenic expression of the F-BAR domain of *Tg*REMIND is toxic for *T. gondii* (17), it was interesting to compare our constructs in a human retinal pigment epithelial-1 (RPE-1) cell line used as a surrogate system. After expressing GFP-*Tg*REMIND for 24 h in the cells, we made two observations by fluorescence microscopy: the protein was mostly cytosolic, and the Golgi apparatus, whose cis- and trans-cisternae were labeled with GM130 and TGN46, respectively, was disorganized and compacted **(Fig. 4A)**. In cells where GFP-BAR_REMIND_ was expressed, we observed even more drastic changes in the Golgi morphology along with the apparition of green puncta dispersed throughout the cytoplasm. In contrast, GFP-REMIND was fully cytosolic, and the Golgi structure was unaltered. Jointly, these results confirm that the BAR but not the REMIND domain confer to *Tg*REMIND the capacity to perturb cell membranes.

**Figure 4.**
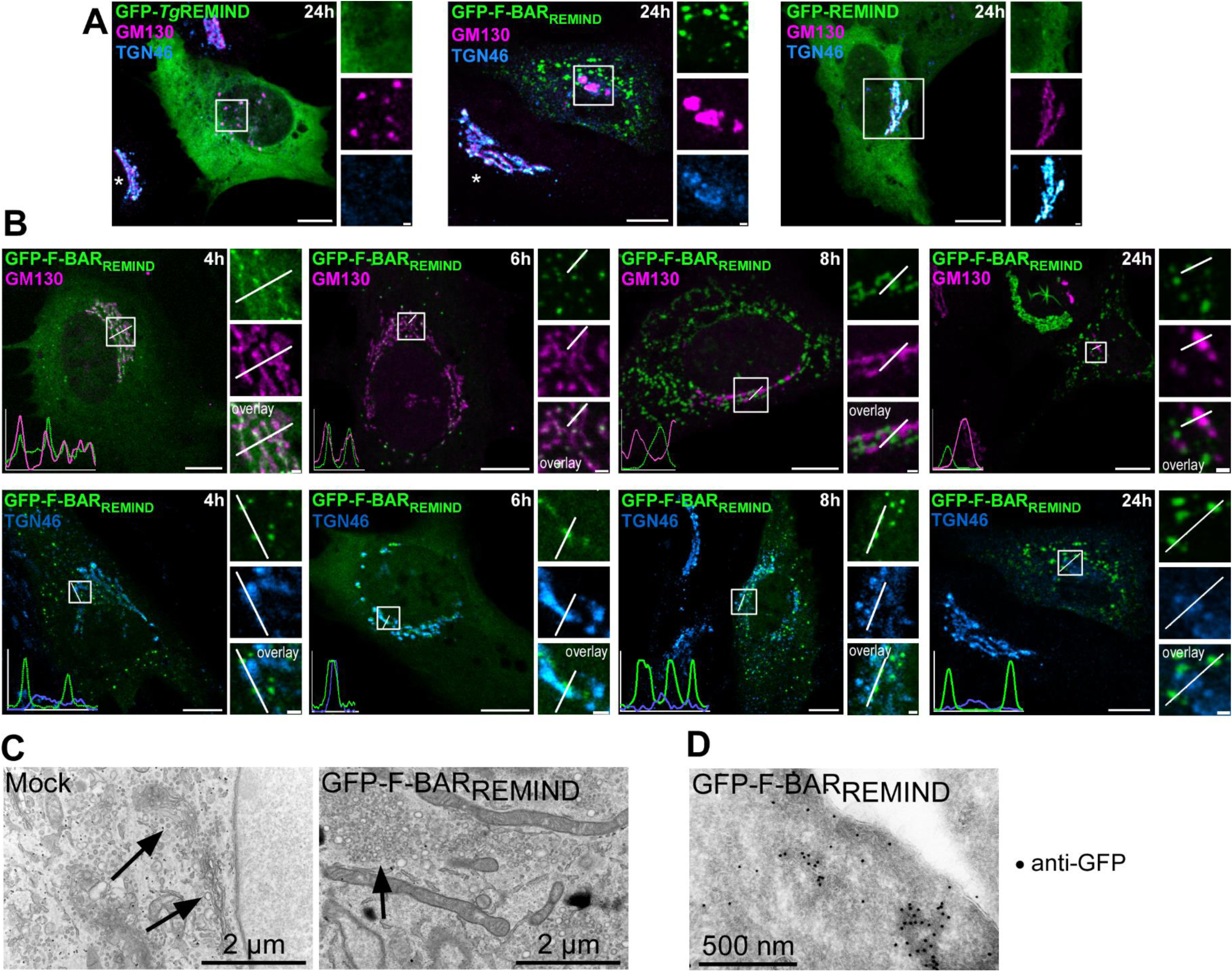
F-BAR_REMIND_ binds to the surface of the Golgi apparatus and causes its disruption. Localization of GFP-F-BAR_REMIND_, GFP-*Tg*REMIND, or GFP-REMIND constructs expressed for 24 h in RPE-1 cells. Before observations by confocal microscopy (Leica TCS SP8, 63×, NA 1.4), cells were fixed and then labeled with an anti-GM130 antibody (magenta) and anti-TGN46 antibody (blue). Stars indicate the presence of a standard Golgi apparatus in non-transfected cells. **(B)** Localization of GFP-BAR_REMIND_ in RPE-1 cells at different time points (4, 6, 8, or 24 h) after transfection. Cells were fixed and labeled with antibodies against GM130 or TGN46. The overlay panel shows merged channels. Line scan shows fluorescence intensities of the green, magenta, or blue channels along the white arrows shown in the insets. Scale bars = 10 µm (inset, 1 µm). **(C)** EM images of RPE-1 cells transfected for 24 h with empty vector (mock) or vector expressing GFP-BAR_REMIND_. The arrow points to a standard Golgi apparatus (left picture) or vesicle clusters (right image). Scale bar = 2 µm. **(D)** Immunogold labeling. GFP-BAR_REMIND_ (beads: 5 nm) are enriched in aggregates localized in the cytosol. Scale bar = 500 nm.

To better understand the properties of F-BAR_REMIND_, we analyzed its cellular localization and the structure of the Golgi apparatus at different time points after transfection of RPE-1 cells (4, 6, 8, and 24 h). We observed that the ribbon-like structure of the Golgi apparatus was maintained at short times (≤ 6 h) and that the GFP-F-BAR_REMIND_ signal was co-localized or juxtaposed to that of GM130 and TGN46, which suggested that the BAR domain was bound to the Golgi membrane **(Fig. 4B)**. We confirmed that, after a longer post-transfection time, the Golgi apparatus underwent intense disorganization and that many dispersed green dots appeared in the cytoplasm. To better understand this phenotype, we analyzed the ultra-structure of cells in which GFP-F-BAR_REMIND_ was expressed for 24 h, by transmission EM. Compared to cells transfected with an empty plasmid (mock), we observed clusters of small vesicles in place of the typical Golgi structure, indicative of its fragmentation, in line with our observations by light microscopy **(Fig. 4C)**. Additionally, using immunogold labeling we identified aggregates that could be strongly labeled with an anti-GFP antibody, indicating that they were composed of F-BAR_REMIND_. Because the expression of F-BAR_REMIND_ did not perturb the cytoskeleton organization, we concluded that this construct likely altered the Golgi structure by directly interacting with its surface **(Fig. S4A)**. In cells expressing *Tg*REMIND, we observed a gradual disorganization of the Golgi apparatus over time but not a clear co-localization or proximity between this construct and Golgi markers **(Fig. S4B)**. As expected, no change in the Golgi structure was seen with GFP-REMIND **(Fig. S4B)**. These results support the idea that the BAR domain but not the REMIND can associate with membranes, leading to their morphological change, corroborating our *in vitro* data. Interestingly, *Tg*REMIND localized weakly with Golgi markers compared to its BAR domain alone, suggesting that the REMIND domain can somehow modulate the association of the F-BAR_REMIND_ domain with membranes.

### REMIND domain prevents *Tg*REMIND from interacting with membranes *via* its F-BAR domain

To further define the potential role of the REMIND domain, we first examined how the full-length *Tg*REMIND bound to membranes compared to F-BAR_REMIND_. Using flotation assays, we found that it was weakly associated with PI(4,5)P_2_-containing liposomes, even those of the smallest size (≤ 30% of total protein), comparatively to F-BAR_REMIND_ **(Fig. 5A**, also compare to data shown in **Fig. 1 and 2).** This supported the idea that the REMIND domain could regulate the membrane-binding capacity of the F-BAR_REMIND_ domain. To explore this, we performed flotation assays in which F-BAR_REMIND_ was incubated with liposomes in the absence or presence of a stoichiometric amount of REMIND. Remarkably, we found that the percentage fraction of F-BAR_REMIND_ bound to small liposomes composed of PC dropped from ∼44 to 8% if it was premixed with REMIND **(Fig. 5B)**. Similar results were obtained with liposomes doped with 10% PI(4,5)P_2_. To confirm these results, we evaluated whether F-BAR_REMIND_ could tubulate liposomes in the presence of the REMIND domain by negative staining EM. Contrary to the control experiment performed with the F-BAR domain alone, we observed no tubulation despite an extensive inspection **(Fig. 5C)**. Collectively, these data strongly suggest that the REMIND domain can prevent the F-BAR_REMIND_ domain from binding membranes.

**Figure 5.**
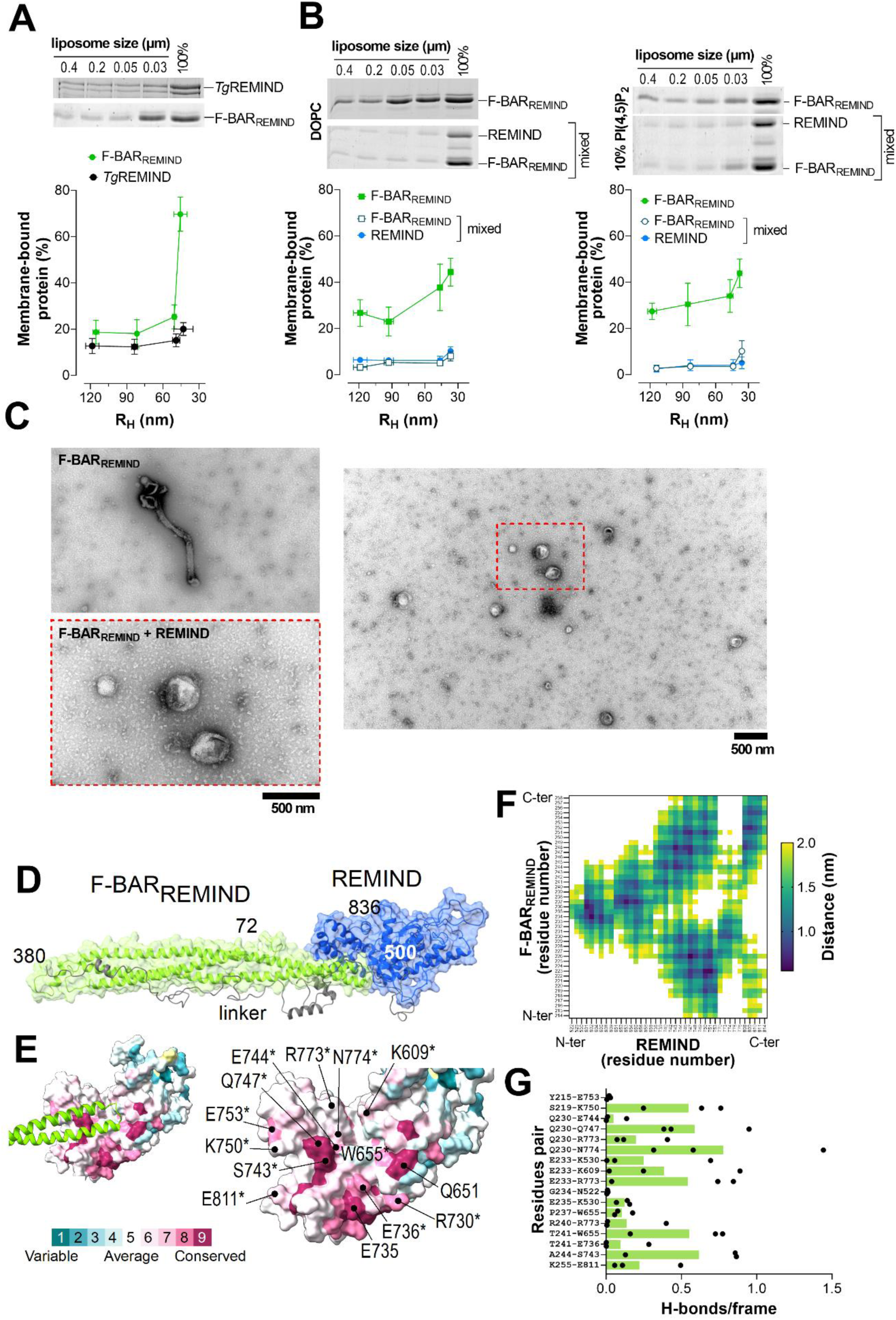
REMIND-dependent inhibition of the membrane-binding capacity of *Tg*REMIND. **(A)** Flotation assays. *Tg*REMIND (0.75 µM) was incubated with liposomes (750 µM lipids) of different sizes, composed of DOPC/PI(4,5)P_2_ (90:10) for 1 h at 25 °C under agitation. Data are represented as mean ± s.e.m. (n = 3). **(B)** F-BAR_REMIND_ was mixed alone or together with a stoichiometric amount of REMIND with liposomes of different sizes, composed of DOPC only or DOPC/PI(4,5)P_2_ (90:10) for 1 h at 25 °C under agitation. Mean ± s.e.m. (n = 3-5). **(C)** Negative staining EM. Representative images of Folch fraction liposomes incubated with F-BAR_REMIND_ alone or mixed with REMIND. A large view shows that no tubule emanates from liposomes when the REMIND domain is present. **(D)** A three- dimensional model of the full-length *Tg*REMIND established by AlphaFold shows the association between the REMIND domain and the tip of the F-BAR domain. Only one monomer is shown. **(E)** Close-up view of F-BAR binding site predicted at the surface of the REMIND domain showing the degree of amino acid conservation based on 37 distinct sequences from diverse Apicomplexan species. Residues that are highly conserved and/or able to form one or more hydrogen bonds with residues of the BAR domain are indicated (a star indicates whether a residue forms hydrogen bond(s)). **(F)** A heat map based on a proximity matrix shows that the 214-258 region of *Tg*REMIND (the extremity of the F-BAR domain) is closely associated with different residues of the REMIND domain. The average distance between two residues was calculated based on conformations observed in one 250-ns MD trajectory. Values higher than 2 nanometers are not shown. **(G)** The average number of H-bonds *per* configuration between two given residues belonging to the BAR domain and the REMIND domain was calculated from three independent 250-ns MD trajectories (green bars). The values obtained independently from each trajectory are also shown (black dots).

Remarkably in line with our data, the analysis of structural models of the *Tg*REMIND protein (encompassing residues 72 to 838 of the full-length protein) predicted that the REMIND domain is associated with the tip of the F-BAR_REMIND_ domain (**Fig. 5D, E**, and (17)). We ran three independent 250 ns-long MD simulations of the protein in solution to further analyze this potential intramolecular interaction. We observed a high increase in the RMSD values **(Fig. S5C)**, which mainly corresponds to significant movements of the linker region between the BAR and REMIND domains and some fluctuations inside the REMIND domain, as indicated by visualizing the MD trajectories (see **Sup. Movie 1)** but also by calculating the RMSF for each residue **(Fig. S5D)**. Along with this, we observed that the extremity of the BAR domain (region 214-259) remained in close association with the REMIND domain along each trajectory, as indicated by the average distances and associated standard-deviation values calculated for pairs of residues, one belonging to the F-BAR domain, the other to the REMIND domain **(Fig.5F, Fig.S6)**. Interestingly, a search for evolutionary conservation in *Tg*REMIND sequences revealed that the region of REMIND in interaction with the F-BAR domain includes many highly conserved residues **(Fig.5 E-F, Fig. S6E, S7)**. Analysis of MD trajectories showed that some of these (K530, K609, E736, S743, E744, Q747, E753, R773, N774, E811, conservation score ≥ 7) can form one or more H-bonds with residues belonging to the F-BAR domain, some of these being also well-conserved (Y215, S219, Q230, P237, R240, T241, A244, **Fig. 5 E-G, Fig. S7**). These data support the notion that the REMIND domain can stably interact with the BAR domain to prevent its interaction with membranes.

### *Tg*BAR2 strongly binds and deforms anionic membranes

A second protein in *T. gondii* of unknown function (GenID TGME49_320760) composed of 357 amino-acids was predicted by AlphaFold to contain a BAR domain (region 14-213) followed by a disordered region **(Fig. S6A, Fig. S8A)**. A close analysis of the structural model suggested that the BAR domain may be adapted to highly-curved membranes, able to fit a membrane tube of ∼22 nm in diameter. Moreover, we estimated that its concave face is extremely positively charged compared to that of F- BAR_REMIND_ **(Fig. 6B)** but also even more basic than that of archetypical BAR and N-BAR domains belonging to amphiphysin, endophilin, and arfaptin **(Fig. S8B)**. To define the membrane-binding properties of this protein (hereafter called *Tg*BAR2), we expressed its recombinant form in *E. coli* and purified it by affinity chromatography and SEC. However, the purity of the preparation was not high enough to accurately assess the folding of the protein by CD spectroscopy. Therefore, we directly analyzed its ability to associate with liposomes of given compositions and sizes using flotation assays. We found that the protein was hardly associated with DOPC liposomes, whatever their size, but, in contrast, was firmly bound to membrane enriched with either 10% PI(4,5)P_2_ or 30% DOPS in a somewhat curvature-dependent manner **(Fig. 6C)**. Next, we directly compared the membrane-binding properties of *Tg*BAR2 with those of F-BAR_REMIND_ in the same flotation assays. We confirmed that F- BAR_REMIND_ but not *Tg*BAR2 was associated with pure PC membranes. An opposite result was obtained with membranes enriched in PS and additionally in PI(4,5)P_2_ **(Fig. 6D).** Next, we investigated whether *Tg*BAR2 could remodel liposomes composed of Folch fraction I lipids at P/L = 1/15 by negative-staining EM. Remarkably, for each condition, we found that *Tg*BAR2 caused the massive formation of narrow and wider tubes with a mean diameter of 16.0 ± 0.5 and 33.6 ± 0.6 nm, respectively **(Fig. 6E, F)**, *i.e.,* narrower than those generated by F-BAR_REMIND._ Moreover, these tubules were, on average, longer than those generated by F-BAR_REMIND_ (1117 ± 89.4 nm as measured for the ticker tubules, **Fig. 6F**). It is also noteworthy that *Tg*BAR2 generated more tubules *per* liposome than F-BAR_REMIND_ **(Fig. S8C)**. To get a higher-resolution view of the membrane-remodeling capacity of *Tg*BAR2, we examined how the protein deformed liposomes by cryo-EM. We confirmed its ability to induce the formation of narrow and broader tubules **(Fig. 6G, H)**. More precisely, we found that the protein transformed liposomes into tubular micelles (**Fig. 6G**, black arrow) and bilayered tubules (white arrow). *Tg*BAR2 coated the membrane surfaces and interacts with the leaflets. Transmembrane densities were also visible (**Fig. 6G**, yellow arrows). By analyzing the sequence of *Tg*BAR2, we did not identify any membrane-binding amphipathic helices or hydrophobic wedges that might account for the remodeling capacity of this protein **(Fig. S8D)**. Collectively, these data suggest that the structural and electrostatic features of the BAR domain of *Tg*BAR2 give this protein a much higher propensity to bind and deform negatively charged membranes compared to F-BAR_REMIND_.

**Figure 6.**
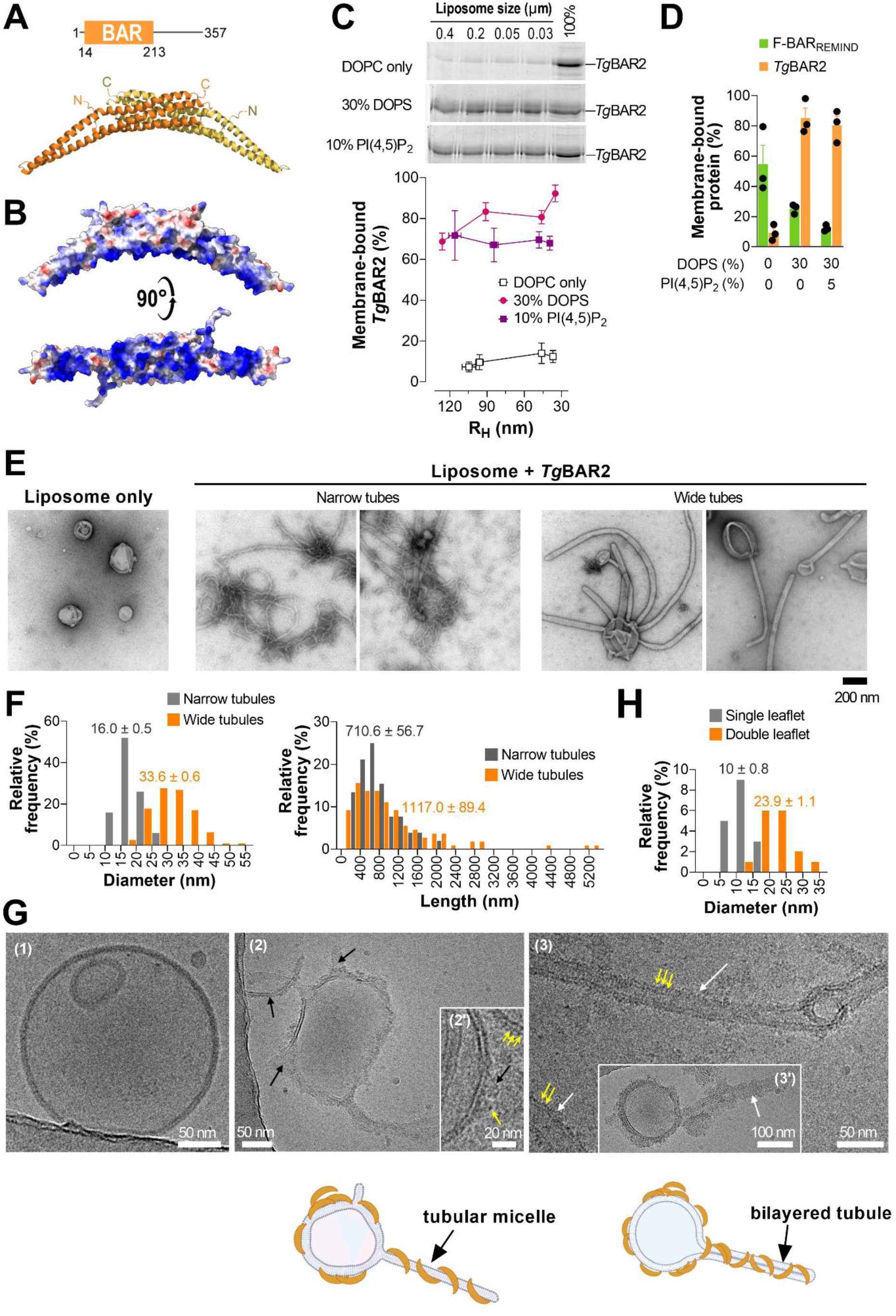
Membrane-binding properties of a second *T.gondii* BAR-domain-containing protein. **(A)** Structural organization of *Tg*BAR2 and AlphaFold-predicted model of its BAR domain in a dimeric form. Its intrinsic curvature seems adapted to the recognition of 22 nm diameter tubules. The position of the N- and C-terminal ends of each monomer is shown. The structure is represented in ribbon mode. **(B)** Electrostatic potential of the dimeric BAR domain red (red = -16.9 kTe^-1^, blue = +16.9 kTe^-1^). **(C)** Flotation assays. *Tg*BAR2 (0.75 µM) was incubated with liposomes of different radii (750 µM lipids), only made of DOPC or additionally containing 30% DOPS or 10% PI(4,5)P_2_, in TN buffer for 1 h at 25 °C under agitation. The percentage of membrane-bound *Tg*BAR2 is represented as the function of the average hydrodynamic radius (R_H_) of liposomes. Data corresponds to mean ± s.e.m. (n = 3). **(D)** Flotation assays. Membrane-bound fraction of *Tg*BAR2 or F-BAR_REMIND_ (0.75 µM) incubated with liposomes extruded through 0.1 µm pores (750 µM lipids), only made of DOPC or additionally containing 30% DOPS and 5% PI(4,5)P_2_. Data corresponds to mean ± s.e.m. (n = 3). **(E)** Negative-staining EM. Liposomes made of Folch fraction I lipids (30 µM) and extruded through 0.4 µm pores were incubated with *Tg*BAR2 (1.9 µM). Control experiments were conducted with liposomes only. Representative pictures are shown. Scale bar = 200 nm. **(F)** Diameter and length distribution of narrow and broader membrane tubules induced by *Tg*BAR2 (narrow tubules, n = 50; wide tubules, n = 112). The indicated values correspond to mean ± s.e.m. **(G)** Cryo-EM. Liposomes made of Folch fraction I lipids (150 µM lipids) and extruded through 0.4 µm pores were mixed with *Tg*BAR2 (5 µM) at P/L= 1/30 and dialyzed three times under agitation in TN buffer for 30 min at cold temperature. A control picture showing liposomes without *Tg*BAR2 is shown (1). *Tg*BAR2 can transform liposomes into tubular micelles (**G,** picture 2 and 2’ black arrow) and bilayered tubules (**G**, picture 3 and 3’, white arrow). *Tg*BAR2 coats the membrane surfaces and can form transmembrane densities (yellow arrows). (G) Lower panels are general interpretations of membrane destabilization phenomena by *Tg*BAR2; left: tubular micelles; right: bilayered tubules **(H)** Diameter distribution of membrane tubules with average values observed by cryo-EM. The indicated value corresponds to mean ± s.e.m. (tubular micelle, n = 16; bilayered tubules, n = 17).

## Discussion

The formation of specific secretory compartments in Apicomplexans, containing many of the factors the parasite uses to invade and reside in its host, relies on specialized vesicular trafficking pathways, the features of which remain poorly described. In this context, the role of BAR-domain-containing proteins is barely known. This study reports the first biochemical characterization of *Tg*REMIND and *Tg*BAR2, two BAR-containing proteins expressed in *T. gondii*, a model parasite of the Apicomplexan phylum.

First, we demonstrate that the N-terminal region of *Tg*REMIND contains an F-BAR domain (F- BAR_REMIND_). We show that this region is well-folded, has a CD signature that matches the structural predictions, and has a dimerization capacity. Furthermore, we establish that this domain associates with liposomes less than 60 nm in radius better than with liposomes of higher radius, indicating that it is adapted to positively-curved membranes, as observed for some BAR domains [*e.g.,* in centaurin (21)) and F-BAR domains [*e.g.,* in syndapin 1)(33)]. Most notably, using a gold-standard procedure based on the observation of liposomes by negative-staining EM (21,29), we find that F-BAR_REMIND_ can generate ∼ 40 nm diameter tubules. Remarkably, as estimated from our structural model, the intrinsic curvature of F-BAR_REMIND_ is perfectly compatible with generating such tubules. Knowing that F-BAR domain-containing proteins can form tubules with diameter > 25-30 nm [*e.g*., F-BAR domain of FCho2 (29,30), Syndapin/Pacsin (25,30,34), and CIP4 (27,30)] whereas proteins with BAR and N-BAR domains predominantly generate narrower tubules ≤ 30 nm [N-BAR domain of amphiphysin and endophilin-A, BAR domain of arfaptin (21,35,36)], our data demonstrates that *Tg*REMIND contains a *bona fide* F-BAR domain. It is worth noting that many F-BAR-containing proteins deform liposomes into a wide range of tubule diameters due to their capacity to adopt diverse orientations relative to the tubule axis (27,29,37).In our assays, F-BAR_REMIND_ generates tubules that are relatively homogenous in diameter, suggesting that this domain adopts one major single orientation at the membrane surface.

F-BAR_REMIND_ can substantially bind to pure PC membranes, and this association is only slightly enhanced by PIPs, whereas an abundance of PS inhibits it. This indicates that *Tg*REMIND has membrane-binding specificities that differ from those of many F-BAR-containing proteins (*e.g.,* FBP17, pacsin/syndapin, FER, PSTPIP) that scarcely associate with membranes unless these contain PS and PIPs (30,31,38). Instead, F-BAR_REMIND_ has some resemblances with the F-BAR domains of yeast proteins Hof1p, Bzz1p, and Rdg1p (23), which all can significantly associate with PC membranes but not necessarily more with membranes enriched with PS or PI(4,5)P_2_, except Rdg1p. Along with this, we found that introducing anionic residues in place of several basic residues, which compose a noticeable cluster in the concave membrane-binding interface of F-BAR_REMIND_, only slightly decreases the avidity of this domain for PI(4,5)P_2_-rich membranes. Introducing anionic residues in a hydrophobic region close to the basic cluster has much more impact. However, puzzlingly, F-BAR_REMIND_ and its mutated forms can deform liposomes made of Folch fraction lipids*, i.e.,* a mixture containing a high proportion of PS. Likely, the high protein level relative to liposome surface in tubulation assays overcomes the low affinity of F-BAR_REMIND_ for negatively charged membranes and promotes binding and bending processes. We conclude that the F-BAR_REMIND_ domain likely binds to membranes *via* hydrophobic and short-range electrostatic interaction rather than long-range electrostatic interaction.

*Tg*REMIND is promiscuously located at many internal compartments (ELC, trans-Golgi, dense granules, and rhoptries (17). We have some clues that the membrane of rhoptries is very rich in PC and sphingomyelin (39) but poor in PI and PS. On the other hand, a recent study has suggested that the Golgi/*trans*-Golgi compartments, but not necessarily dense granules and rhoptries, contain a PI(4)P pool (40). However, despite these differences, *Tg*REMIND is localized on all these subcellular compartments, suggesting a capacity to target these compartments via non-specific hydrophobic interaction in a PIP-independent manner. In agreement with this, we observed that F-BAR_REMIND_, when expressed in RPE-1 cells, is primarily found at the *cis*-Golgi level, which is known to be poor in negatively charged lipids and devoid of PIPs. Finally, Wan et al. also observed that *Tg*REMIND (called BAR1) is localized on internal compartments in *T.gondii* but is neither present at the PM nor co- localized with GAP45, a marker of the medium IMC (16). Knowing that the inner leaflet of the medium IMC and PM is rich in PS (41), this observation fits well with our findings that the F-BAR_REMIND_ barely associates with PS-rich membranes *in vitro*. Moreover, *Tg*REMIND is not localized at the PM where PI(4,5)P_2_ is prominent (40), suggesting that this lipid is not a targeting determinant for that protein. Collectively, given that the REMIND domain has no membrane-binding capacity (see discussion below), we suggest that the F-BAR domain of *Tg*REMIND, as it can bind neutral membranes with low specificity and no strict requirement for PIPs, allowed this protein to be ubiquitously localized on various organelles in *T. gondii*.

In addition to characterizing the membrane-binding properties of F-BAR_REMIND_, our study provides some insights into the potential function of *Tg*REMIND. We observed that F-BAR_REMIND_ tends to oligomerize in solution. However, we did not observe any ability of this domain to form organized structures on tubules, *i.e.,* coat as other F-BAR-containing proteins (27,29). This suggests that its potential role is not to stabilize tubular structures. Moreover, we observed by negative staining that F- BAR_REMIND_ only induced the formation of one, sometimes two, tubules *per* liposome, and we failed to observe tubules by cryo-EM. However, by this approach, we observed that F-BAR_REMIND_ could bind to and disorganize the bilayer structure of liposomes. We also noted that, when overexpressed in RPE-1 cells, F-BAR_REMIND_ did not induce observable membrane tubulation as found for many other BAR and F-BAR domains ((31,32) but disrupted the Golgi apparatus. Therefore, F-BAR_REMIND_ has a dual ability to deform and disrupt membranes. Such a capacity to break membranes could explain why expressing this domain alone is highly toxic for *T. gondii* (17). These data suggest that *Tg*REMIND has the intrinsic ability to deform and /or disrupt endomembranes.

Next, we fully establish the existence of a new structural domain called REMIND, which was predicted to be in the second half of *Tg*REMIND, connected to the F-BAR domain *via* a disordered linker (17). Key evidence is that the C-terminal region of *Tg*REMIND has a secondary structure content compatible with the tridimensional model of the REMIND domain predicted by AlphaFold. Next, we show that the REMIND domain has no membrane-binding capacities, ruling out previous claims based on crude protein-lipid overlay assays and not authentic lipid bilayers (17). Most remarkably, we found that this domain, added *in trans*, prevents F-BAR_REMIND_ from binding to and remodeling membranes. This means that an intramolecular association between F-BAR_REMIND_ and REMIND domains might be the basis of an autoregulatory mechanism able to tune the association of *Tg*REMIND with membranes. Therefore, the REMIND domain would play a role analogous to that of the SH3 domain, which, inside syndapin 1/Pacsin 1 (42), Drosophila Nervous Wreck (Nwk) (43,44) or endophilin (45,46) interact with the F-BAR (or N-BAR domains in the case of endophilin) to limit the association of these proteins with membranes. Our model is in line with several observations: the full-length *Tg*REMIND poorly binds membrane *in vitro* and does not associate with the Golgi membrane as clearly as does the F-BAR_REMIND_ domain alone when expressed in RPE-1 cells. Also, as mentioned before, the transgenic expression of F-BAR_REMIND_ is toxic for *T. gondii*, which indirectly suggests that the membrane-binding capacity of this domain is tightly controlled when associated with the REMIND domain (17). Moreover, structural predictions combined with MD simulations suggest that REMIND stably interacts with the tip of the F- BAR_REMIND_ *via* evolutionarily-conserved residues. Our data suggest that *Tg*REMIND is a membrane-disrupting device regulated by the REMIND domain. Given that this protein potentially interacts with many trafficking factors (17), we posit that one or more protein partners unlock the auto-inhibition state of *Tg*REMIND and trigger its membrane remodeling capacity at different stages of the trafficking pathways. This might explain why the absence of *Tg*REMIND impacts the biogenesis of dense granules and rhoptries (17). Further work is necessary to explore this possibility.

Next, we report that *T. gondii* expresses a second BAR-domain containing protein, *Tg*BAR2, with features that highly contrast with those of the BAR domain of *Tg*REMIND. Its BAR domain is highly basic and strongly binds membranes, but only if those are enriched with negatively charged lipids. *Tg*BAR2 binds to these membranes in a rather curvature-independent manner and powerfully deforms liposomes into tubules that can be extremely narrow (∼10 nm in diameter). These features are similar to those of many other BAR and N-BAR-containing proteins (21,36,47), suggesting that the function of *Tg*BAR2 would be to form and stabilize highly-curved membrane structures. However, we noted that the concave face of the BAR domain of *Tg*BAR2 is far more basic than that of these archetypical examples of the BAR domain. These features might explain the capacity of *Tg*BAR2 to powerfully deform membranes despite its lack of AH. We additionally observe by cryo-EM that *Tg*BAR2 can induce the formation of tubules only made of one lipid monolayer, a characteristic that has never been reported for any other BAR domain to our knowledge. Therefore, the BAR domain of *Tg*BAR2 represents a new kind of BAR domain. How this is related to the function of that protein in *T. gondii* remains challenging to say. *Tg*BAR2 (under the name BAR2) has been recently reported to be, contrary to *Tg*REMIND, at the periphery of the cell, colocalizing with IMC. As mentioned before, the inner leaflet of the medium IMC is particularly well-enriched in PS, and the inner leaflet of PM contains both PS and PI(4,5)P_2_ (41). Thus, these observations fit well with our observation that *Tg*BAR2 associates *in vitro* with membranes containing both lipids. However, puzzlingly, despite its localization and ability to remodel negatively-charged membrane, *Tg*BAR2 is not found at the level of the micropore, an endocytic structure derived from the parasite PM bearing dynamin-like protein *Tg*DrpC, *Tg*AP-2 adaptor, and Kelch13 (18). However, it should be noted that other endocytic structures have been observed (48,49) but have yet to be characterized from a molecular point of view. The finding that *Tg*BAR2 has a strong capability of remodeling anionic membranes should motivate further investigations.

## Experimental procedures

### Plasmids

The [70-347] region of *Tg*REMIND (F-BAR_REMIND_) was fused to an N-terminal His-tag and cloned into a pET-M13 plasmid (from S. Tomavo and S. Zinn-Justin, I2BC, France). Five cysteines were replaced by alanine or serine residues (C91S, C160A, C218A, C251A, and C296S mutations) to prevent protein aggregation during purification. Additional mutations in this construct were generated using the QuikChange Lightning Site-Directed Mutagenesis Kit (Agilent technologies). The sequence of the full-length *Tg*REMIND or the [495-840] region of the protein (REMIND region) was cloned in a pGEX-6P-3 plasmid to be expressed in fusion with an N-terminal GST tag (from S. Tomavo and S. Zinn-Justin). The tag and each protein of interest are linked by a sequence (LEVLFQGPLGSGGTGQQMGRDLENLYFQG) containing a Protease Precission cleavage site. Mutations in the REMIND construct were introduced by site-directed mutagenesis using PfuUltra II High-fidelity DNA Polymerase (Agilent technologies). A codon-optimized synthetic *Tg*BAR2 gene was cloned in pET-15b plasmid (ProteoGenix) to express the protein with an N-terminal His-tag. The sequence of the N-BAR domain of human amphiphysin (residues 2–242, N-BAR_Amph_) was cloned into a pGEX-4T-2 plasmid to be expressed in fusion to an N-terminal GST tag (gift from Dr. J.C. Stachowiak, University of Texas, USA).

For cell biology experiments, a codon-optimized synthetic *Tg*REMIND gene was cloned in a pEGFP-C1 expression vector to express the protein in fusion with an N-terminal GFP (Proteogenix). A stop codon was inserted in the coding sequence of *Tg*REMIND to express the [1-382] region of *Tg*REMIND appended with an N-terminal GFP (GFP-F-BAR_REMIND_). Another expression vector, coding for the region [500-842] of *Tg*REMIND in fusion with GFP, was also prepared (GFP-REMIND). The sequences of all these constructs and mutants were checked by DNA sequencing.

### Protein expression and purification

All *T.gondii* proteins were expressed in *E. Coli* (BL21-GOLD(DE3)) competent cells (Stratagene) grown in Luria Bertani Broth (LB) medium at 37 °C until the optical density (OD_600_) of the bacterial suspension reached a value of 0.4 and then incubated at 18 °C until OD_600_ reached 0.8. The expression of F-BAR_REMIND_ and its mutated form was induced with 0.1 mM isopropyl β-D-1- thiogalactopyranoside (IPTG) and conducted overnight at 18 °C under agitation at 200 rpm. For *Tg*BAR2, the production occurred in similar conditions, except that the expression of the protein was induced by 0.2 mM IPTG. The bacteria cells were harvested and re-suspended in cold buffer A (50 mM Tris, pH 8, 150 mM NaCl, 10 mM imidazole, 5% (v/v) glycerol) supplemented with 1 mM PMSF, 10 µM bestatin, 1 µM pepstatin A and cOmplete EDTA-free protease inhibitor tablets (Roche). Cells were lysed using a Cell Disruptor TS SERIES (Constant Systems Ltd.) and the lysate was centrifuged at 186,000 × *g* for 90 min. Then, the supernatant was applied to HisPur™ Cobalt Resin (Thermo Scientific) for 3 h 30. The beads were washed twice with buffer A, then twice with high salt buffer (50 mM Tris, pH 8, 800 mM NaCl, 10 mM imidazole, 5% (v/v) glycerol), and twice again with buffer A. Next, the beads were incubated with 1.5 mL buffer A supplemented with 300 mM imidazole for 10 min to eluate the protein, and this step was repeated four times. The collected fractions were pooled and concentrated in an Amicon filter unit (Millipore) and further purified by size-exclusion chromatography using an XK- 16/70 column packed with Sephacryl S-200 HR pre-equilibrated with buffer B (50 mM Tris, pH 8, 150 mM NaCl, 5% (v/v) glycerol). The pure protein fractions were pooled, concentrated, and supplemented with 5% (v/v) glycerol. For all these proteins, aliquots were prepared, flash-frozen in liquid nitrogen, and stored at –80 °C. The concentration of the proteins was determined by measuring their absorbency at λ = 280 nm or on SDS-PAGE compared to a range of BSA concentrations.

Similar purification procedures were applied for *Tg*REMIND or the REMIND construct except that buffer C (50 mM Tris, pH 8, 150 mM NaCl, 5 mM DTT, 5% (v/v) glycerol) supplemented with 1 mM PMSF, 10 µM bestatin, 1 µM pepstatin A and cOmplete EDTA-free protease inhibitor tablets was used to lyse the bacterial cells. Once separated from cell membranes by centrifugation, the supernatant was applied to Glutathione Sepharose™ 4B (Cytiva) beads for 3 h 30. The beads were washed twice with buffer C, followed by two washing steps with high salt buffer (50 mM Tris, pH 8, 800 mM NaCl, 5 mM DTT, 5% (v/v) glycerol), and then washed twice with buffer C. Proteins were cleaved off the GST bound to resin in the presence of preScission protease (Cytiva) in a 2 mL-volume tube overnight at 4 °C with rocking. The proteins were recovered in the supernatant collected by washing the beads several times with 0.7 mL of buffer C and subjected to mild centrifugation. The protein was concentrated using an Amicon filter unit (Millipore) and further purified by SEC using a Superose® 6 Increase 10/300 GL column equilibrated with buffer C (supplemented with 1 mM EDTA when purifying *Tg*REMIND). The pure protein fractions were pooled, concentrated, and supplemented with 5% (v/v) glycerol to reach 10% glycerol. Each protein was stored as described before, and its concentration was determined by spectrometry or SDS-PAGE analysis.

Amphiphysin N-BAR was expressed according to the protocol reported in (50). Once the OD_600_ of the bacterial suspension reached a value of 0.8, the expression of the protein was induced with 1 mM IPTG for 2 h at 37 °C under agitation of 200 rpm. Bacterial cells were harvested and resuspended with buffer C supplemented with 1 mM PMSF, 10 µM bestatin, 1 µM pepstatin A, and cOmplete EDTA-free protease inhibitor tablets (Roche). Cells were lysed as indicated above, and the supernatant was applied to Glutathione Sepharose^TM^ 4B beads. The beads were washed four times with buffer C, two times with highly salted buffer (50 mM Tris, pH 8, 800 mM NaCl, 2 mM DTT, glycerol 5% (v/v)), and then again four times with buffer C. The protein was cleaved from the GST tag by incubating the beads with human thrombin (Sigma-Aldrich) in the presence of 50 µM CaCl_2_ overnight at 4 °C with rocking. The protein was recovered in the supernatant after four cycles of centrifugation and washing the beads with 0.7 mL of buffer C supplemented with 2 mM EDTA. The protein was concentrated using an Amicon filter unit (Millipore) down to 1 mL. The protein was then supplemented with 5% (v/v) pure glycerol, aliquoted, flash-frozen in liquid nitrogen, and stored at –80 °C. The concentration of the proteins was determined using a BSA assay.

### Lipids

18:1/18:1-PC (1,2-dioleoyl-*sn*-glycero-3-phosphocholine or DOPC), 18:1/18:1-PS (1,2-dioleoyl-*sn*- glycero-3-phospho-L-serine or DOPS), brain PI(4)P (L-α-phosphatidylinositol 4-phosphate), brain PI(4,5)P_2_ (L-α-phosphatidylinositol 4,5-bisphosphate), NBD-PC (1-palmitoyl-2-(12-[(7-nitro-2-1,3- benzoxadiazol-4-yl)amino]dodecanoyl)-sn-glycero-3-phosphocholine) NBD-PE (1,2-dioleoyl-*sn*-glycero-3-phosphoethanolamine-N-(7-nitro-2-1,3-benzoxadiazol-4-yl)) were purchased from Avanti Polar Lipids. Lipids from bovine brain extract (Folch fraction 1) and 16:0/16:0-PI(3)P were purchased from Sigma Aldrich and Echelon Biosciences, respectively.

### Liposomes preparation

Lipids stored in CHCl_3_ or CHCl_3_/methanol stock solutions were mixed at the desired molar ratio. The solvent was removed in a rotary evaporator under vacuum. If the flask contained a mixture with PIPs, it was pre-warmed at 33 °C for 10 min before creating a vacuum. The lipid film was hydrated in 50 mM Tris, pH 7.4, 150 mM NaCl (TN) buffer to obtain a suspension of multi-lamellar vesicles. This suspension was frozen and thawed five times and then extruded sequentially through polycarbonate filters with pore sizes of 0.4, 0.2, 0.1, 0.05, and 0.03 μm. The liposome hydrodynamic radius (R_H_) was estimated by dynamic light scattering using a Dyna Pro instrument. Liposomes were stored at room temperature and in the dark when containing fluorescent lipids and used within two days.

### Flotation assay

Flotation assays using liposomes of different radii were performed as described in (19). Protein association to the membrane was measured by incubating the protein (0.75 µM) with NBD-PC containing liposomes (750 µM lipids) for 1 h at 25 °C. Then, the suspension was adjusted to 28% (w/w) sucrose by mixing 100 µL of a 60% (w/w) sucrose solution in TN buffer and overlaid with 200 µL of TN buffer containing 24% (w/w) sucrose and 50 µL of sucrose-free TN buffer. The sample was centrifuged at 240,000 × *g* in a swing rotor (TLS 55 Beckmann) for 70 min. The bottom (250 µL), middle (140 µL), and top (110 µL) fractions were collected. The bottom and top fractions were analyzed by SDS-PAGE after staining with SYPRO Orange using a FUSION FX fluorescence imaging system.

### Electron microscopy

Liposomes (30 μM total lipids) extruded through 0.4 μm pores were mixed with protein (1.9 μM for condition P/L = 1/15) in TN buffer for 2 h. For negative staining imaging, one drop of ∼10 μL was placed on the top of the copper EM grid covered with a formvar film for 5 min and then dried and stained with uranyl acetate (1% in distilled water). The grid was examined using a JEOL JEM 1400 transmission electron microscope at 100 kV equipped with a MORADA CCD 11 MPixels camera (Olympus SIS).For cryo-electron microscopy, 4 µL of the sample were deposited onto glow-discharged Quantifoil R2/2 holey carbon grids, blotted for 4 s with filter paper, and plunged into liquid ethane using a Vitrobot Mark IV (ThermoFisher Scientific) operated at room temperature and 100% relative humidity. The cryo specimens were transferred into a Gatan 626 cryo-holder and observed in a JEOL 2010F electron microscope operated at 200 kV. Images were recorded at a nominal defocus of 2 μm on a Gatan K2 Summit camera under low electron-dose conditions. Images were denoised by wavelet filtration in ImageJ (https://imagej.net/ij/) using the plugin “A trous filter,” k_1_=20, k_n>1_=0.

### Circular dichroism

The experiments were performed on a Jasco J-815 spectrometer at room temperature with a quartz cell of 0.05 cm path length. Proteins were dialyzed three times against 20 mM Tris, pH 7.4, 120 mM NaF buffer for 30 min to remove glycerol or DTT contained in protein stocks and to exchange buffer. Each spectrum is the average of ten scans recorded from λ = 190 to 260 nm with a bandwidth of 1 nm, a step size of 0.5 nm, and a scan speed of 50 nm.min^-1^. Protein concentration was determined at λ = 280 nm by spectrometry or by densitometry in SDS-PAGE against a BSA concentration range. A control spectrum of buffer was subtracted from each protein spectrum. The percentages of protein secondary structure were estimated by analyzing their CD spectrum (in the 190-250 nm range) using the BeStSel method provided online (51) and compared to the percentages derived from the analysis of Alphafold- predicted structural model and conformations adopted by this model during MD simulations.

### Analytical gel filtration

The various constructs were analyzed by gel filtration on a Superose® 6 Increase 10/300 GL column equilibrated in TN buffer (50 mM Tris, pH 7.4, 150 mM NaCl). Calibration was performed using the following standards: apoferritin (molecular weight 443 kDa, Stokes radius 6.1 nm), alcohol dehydrogenase (150 kDa, 4.5 nm), bovine serum albumin (67 kDa, 3.6 nm), carbonic anhydrase (31 kDa, 2.4 nm) and cytochrome c (12.6 kDa, 1.6 nm).

### Tryptophan-fluorescence assay

Emission fluorescence spectra of protein (1 μM) in TN buffer were measured at 30 °C from 300 to 450 nm (excitation at λ_ex_ = 280 nm) in a quartz cell. The fluorescence of L-tryptophan (zwitterionic form) was recorded as a comparison.

### Structural analysis

The electrostatic surfaces of proteins were calculated using ChimeraX software (52) with the Coulombic electrostatic potential (ESP).

### Sequence analysis

To analyze the evolutionary conservation of amino acid positions in *Tg*REMIND, protein sequences were downloaded from UniProt and analyzed with BLAST (Basic Local Alignment Search Tool) at NCBI (National Center for Biotechnology Information) using a non-redundant protein sequence database from which proteins from *T. gondii* were excluded. Alignments were analyzed with UGENE (53). The phylogenetic tree was visualized using the ITOL website (54). Based on the generated multiple sequence alignment (MSA), the ConSurf server was used to analyze the evolutionary conservation of amino acid positions (55–57), and the results were displayed on the three-dimensional model of *Tg*REMIND using ChimeraX software. Heliquest web server (58) was used to calculate the mean hydrophobicity, mean hydrophobic moment, and net charge along the sequence of *Tg*REMIND and *Tg*BAR2 using an 18-aa overlapping window.

### Molecular dynamics simulations

The tridimensional structure of *Tg*REMIND (segment 72-836) has been predicted using AlphaFold (17,59). Only the structure of the 500-836 region was used for the MD simulation of the REMIND domain. In all cases, the system was built with the protein or domain immersed in a water box using the TIP3P water model and was minimized and equilibrated with 120 mM NaCl using GROMACS 2021.4 Molecular Dynamics (MD) simulation engine (60) using the CHARMM36m force field (61) following the Charmm-Gui procedure (62). MD simulations were performed with GROMACS 2021.4. Bonds involving hydrogen atoms were constrained using the LINCS algorithm (63), and the integration time step was set to 2 fs. The V-Rescale thermostat (64) was used to keep a temperature at 310 °K with a coupling time constant of 1 ps. For simulations with constant pressure, the Parrinello-Rahman barostat (65) was used to maintain a pressure of 1 bar with a compressibility of 4.5 × 10^-5^ and a coupling time constant of 2 ps. Van der Waals (VDW) interactions were switched to zero over 10 to 12 Å, and electrostatic interactions were evaluated using the Particle Mesh Ewald (PME) method (66). For *Tg*REMIND, three different MD simulations of 250 ns-length were launched using different initial seed velocities. For the REMIND domain, a 1 µs-length simulation was launched.

### Analysis of MD simulations

Root Mean Square Deviation (RMSD), Root Mean Square Fluctuation (RMSF), H-bond analysis, and distance measurements between residues (considering mass center) were carried out using GROMACS 2021.4 (60). Home-made scripts based on Python were used to build the distance matrix. All molecular pictures and movies were made with Pymol and Visual Molecular Dynamics (VMD) (67). Circular dichroism spectra were predicted from MD-generated REMIND structure using PDBMD2CD (68).

### Cell culture and transfection

hRPE-1 cells (hTERT-immortalized retinal pigment epithelial cells) were obtained at ATCC (CRL- 4000). Cells were maintained and grown in Dulbecco’s modified Eagle medium DMEM/F12 (Gibco) supplemented with 10% fetal calf serum (FCS) and 1% antibiotics (Zell Shield®) at 37 °C under 5% CO_2_. Following the initial growth phase, the cells were seeded at a density of 40,000 cells per condition in an 8-well coverslip (Ibidi 80826) and allowed to grow for 24 h. Subsequently, the cells were transiently transfected with 250 ng of plasmid DNA using the Lipofectamine 3000 transfection reagent (Thermo Fisher Scientific), following the manufacturer’s protocol.

### Immunolabeling, fluorescence microscopy, and image analysis

At different times after transfection, the cells were fixed with paraformaldehyde 4% in PBS (PFA, Fisher) for 15 min. Next, the cells were treated for 30 min with PBS supplemented with BSA 0.2% (v/v) and saponin 0.05% (v/v) at room temperature. The cells were then incubated for 1 h with primary antibodies directed against the *cis*-Golgi marker GM130 (mouse anti-GM130, BD bioscience, 610822) and/or *trans-*Golgi marker TGN46 (sheep anti-TGN46, Bio-Rad, AHP500G). Then, the cells were washed three times with PBS supplemented with BSA and saponin and incubated for 1 h with secondary fluorescent antibodies, a goat anti-mouse antibody Alexa 640, and/or a donkey anti-sheep Alexa 594 (Invitrogen). Cells were rinsed with PBS and then observed using a Leica TCS SP8 STED 3X in confocal mode. Images were analyzed using ImageJ software.

### Electron microscopy of cells

RPE-1 cells transfected with GFP-F-BAR_REMIND_ were fixed with 4% paraformaldehyde and 0.1% glutaraldehyde in 0.1 M phosphate, pH 7.4 buffer for 2 h. They were processed for ultra-cryo-microtomy using a slightly modified Tokuyasu method (69). The cell suspension was spun down in 10% gelatin. After that, the cells were immersed in 2.3 M sucrose in 0.1 M phosphate, pH 7.4 buffer overnight at 4 °C, and then rapidly frozen in liquid nitrogen. Ultrathin (70 nm thick) cryo-sections were prepared using an ultra-cryo microtome (Leica EMFCS, Austria) and mounted on formvar-coated nickel grids (Electron Microscopy Sciences, USA). The grids were incubated successively in PBS containing the relevant primary antibody for 1 h and then incubated with PBS containing 15 nm colloidal gold-conjugated protein AG (CMC, University Medical Center, Utrecht, The Netherlands). Finally, the samples were fixed for 10 min with 1% glutaraldehyde, contrasted with a mixture of methylcellulose/sucrose and 0.3% uranyl acetate on ice. After being dried in air, sections were examined under a JEOL 1400 transmission electron microscope. The gold particles appear as dark spots, indicating the location of the GFP-F- BAR_REMIND_ within the RPE-1 cell.

## Acknowledgments

We thank Dr. J.C. Stachowiak and Drs. S. Zinn and S. Tomavo for plasmids and initial purification protocols. NA and this work were supported by an ANR-19-CE44-0006 grant from the Agence Nationale de la Recherche. The authors acknowledge the Electron Microscopy facility CCMA (Centre Commun de Microscopie Appliquée) from the « Université Côte d’Azur », part of the « Microscopie Imagerie Côte d’Azur » GIS IBiSA labeled platform, supported by Université Côte d’Azur, the “Région Sud » and the Département 06. The authors also acknowledge financial support from the CNRS-CEA network for transmission electron microscopy and atom probe studies (METSA, FR CNRS 3507) on the LPS cryo-EM platform. This work was supported by State aid under France 2030 (PhOM -Graduate School Physique) with reference ANR-11-IDEX-0003.

## Author contributions

G.D. designed and supervised research. N.A. carried out site-directed mutagenesis, produced and purified all the recombinant proteins of this study, performed *in vitro* experiments, and analyzed protein sequence and tridimensional models. S.P. and N.A. conducted negative-staining EM. M.M. and S.L-G conducted cell biology experiments using light and electron microscopy. A.L., J.D., and A.A.A. performed cryo-electron microscopy with N.A. R.G. ran MD simulations. All the authors analyzed the data. G.D. wrote the manuscript. All of the authors discussed the results and commented on the manuscript.

## Data Availability

Raw data and source code are available from the corresponding author(s) upon reasonable request.

## Competing interests

The authors declare no competing interests.

**Supplementary Movie 1.** One example of an all-atom MD simulation of *Tg*REMIND[72 to 838]. The length of the simulation is 250 ns. The protein is represented in a ribbon mode with the α-helices in purple, the 3_10_ helices in blue, and the β-strands in yellow.

---

**Supplementary Figure 1.**
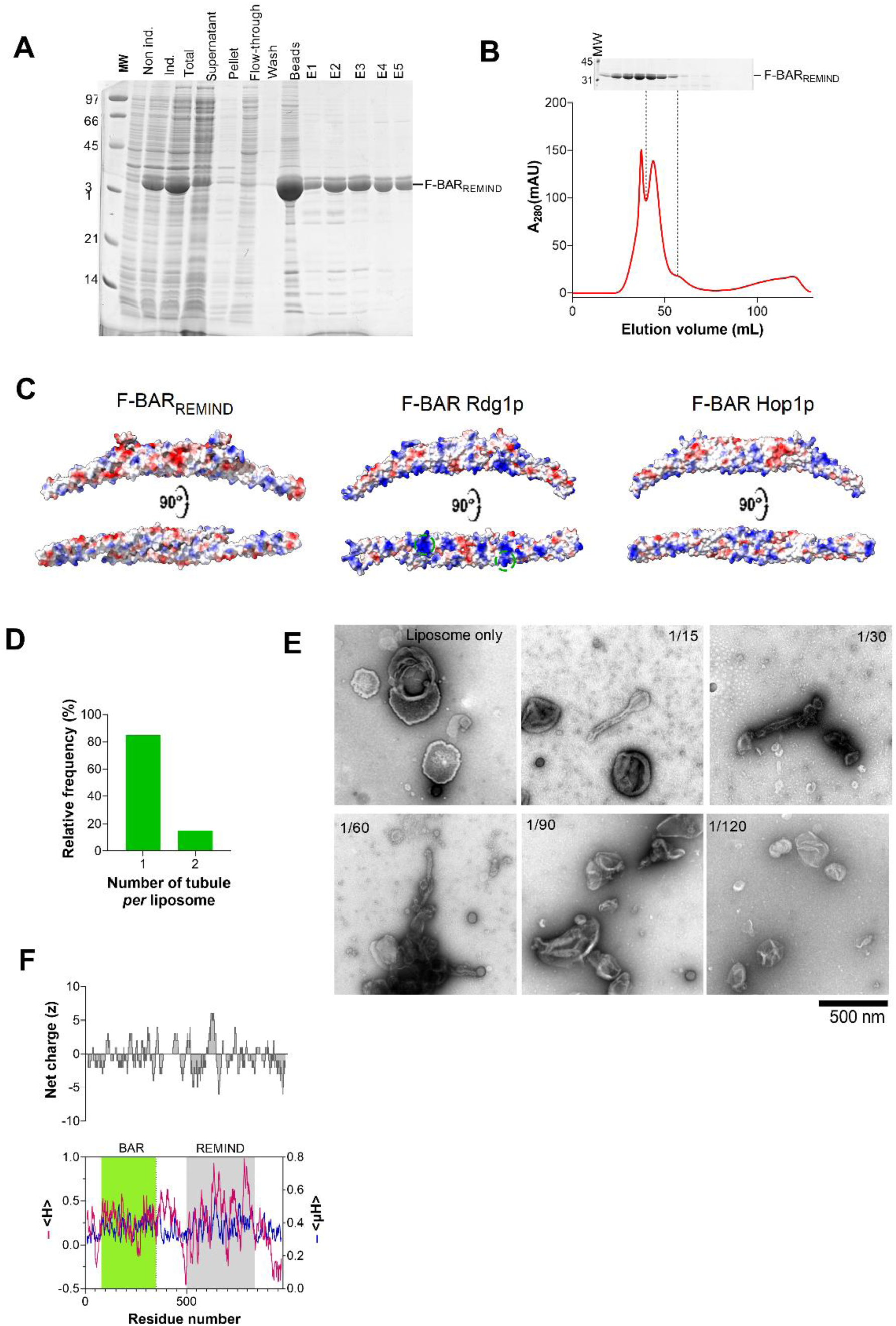
Purification of F-BAR_REMIND_. SDS-PAGE analysis was used to check the presence of the F-BAR_REMIND_ construct at different steps of the purification procedure. MW: molecular-weight size marker; Non ind.: no protein expression induction; Ind.: Induction. E: elution. (see details in Experimental procedures). **(B)** Elution profile of F- BAR_REMIND_ during its purification by SEC. AU: arbitrary unit. The collected fractions have been analyzed by SDS-PAGE to assess the purity of the protein and pooled. **(C)** Electrostatic potential of the dimeric F-BAR_REMIND_ domain and the F-BAR domain of Rdg1p (PDB ID: 4WPC) and Hop1p (4WPE) (red = -16.9 kTe^-1^, blue = +16.9 kTe^-1^). **(D)** Number of tubules *per* liposomes (considering only deformed liposomes, n = 40). **(E)** Negative-staining EM. Liposomes made of Folch fraction I lipids (30 µM lipids) and extruded through 0.4 µm pores were incubated with different concentrations of F- BAR_REMIND_. A control picture of liposomes alone is shown. Scale bar = 500 nm. **(F)** Analysis of the *Tg*REMIND sequence shows that the linker region does not contain any 18-aa segment combining a high mean hydrophobicity (<H>), high hydrophobic moment (<µH>), and net positive charge (z) able to fold into a membrane-binding amphipathic helix. The boundaries of the BAR and REMIND domains are indicated.

**Supplementary Figure 2.**
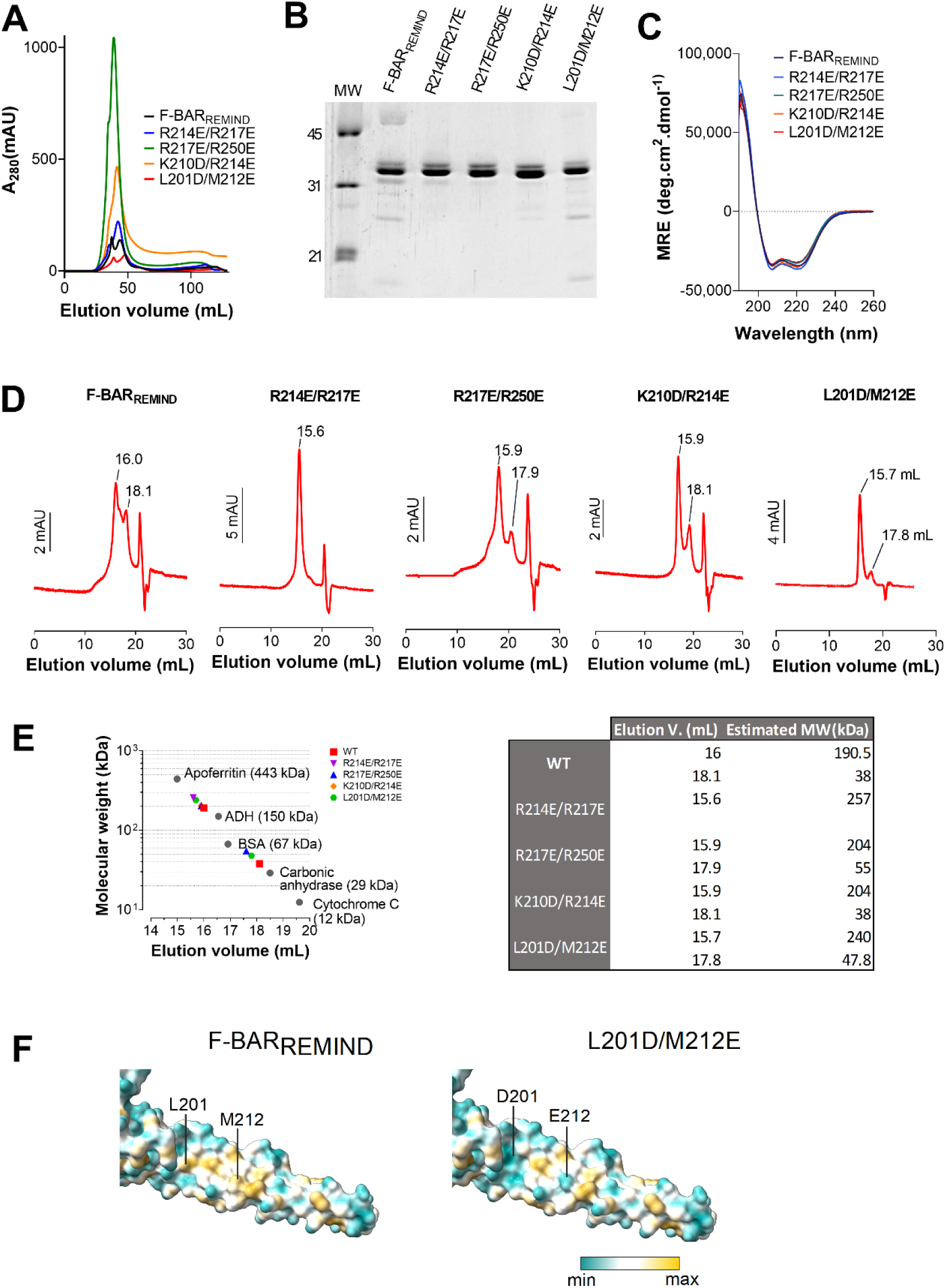
Purification of mutated forms of F-BAR_REMIND_. Elution profile of F-BAR_REMIND_ and R214E/R217E, R217E/R250E, K210D/R214E, and L201D/M212E mutants during the purification of these constructs by SEC. AU: arbitrary unit. **(B)** SDS-PAGE analysis of F-BAR_REMIND_ and its mutated forms showing the purity of each protein batch. **(C)** Far-UV CD spectrum of purified F-BAR_REMIND_ and each mutant (2-10 µM) in 20 mM Tris, pH 7.4, 120 mM NaF buffer recorded at room temperature. MRE: mean residue ellipticity. **(D)** Elution profiles obtained by analytical SEC of F-BAR_REMIND_ and its mutated forms on a Superose 6 Increase 10/300 GL column. The elution volume corresponding to the main peaks is indicated. AU: arbitrary unit. **(E)** Analytical SEC of the constructs. The straight line represents the best fit for the elution of molecular weight standards. **(F)** Close-up view of the lateral side of one F-BAR_REMIND_ monomer in which the L201 and M212 residues have been mutated into D and E residues, respectively. The surface of the wild-type and mutated form of the domain is colored according to the hydrophobicity of the residues using ChimeraX.

**Supplementary Figure 3.**
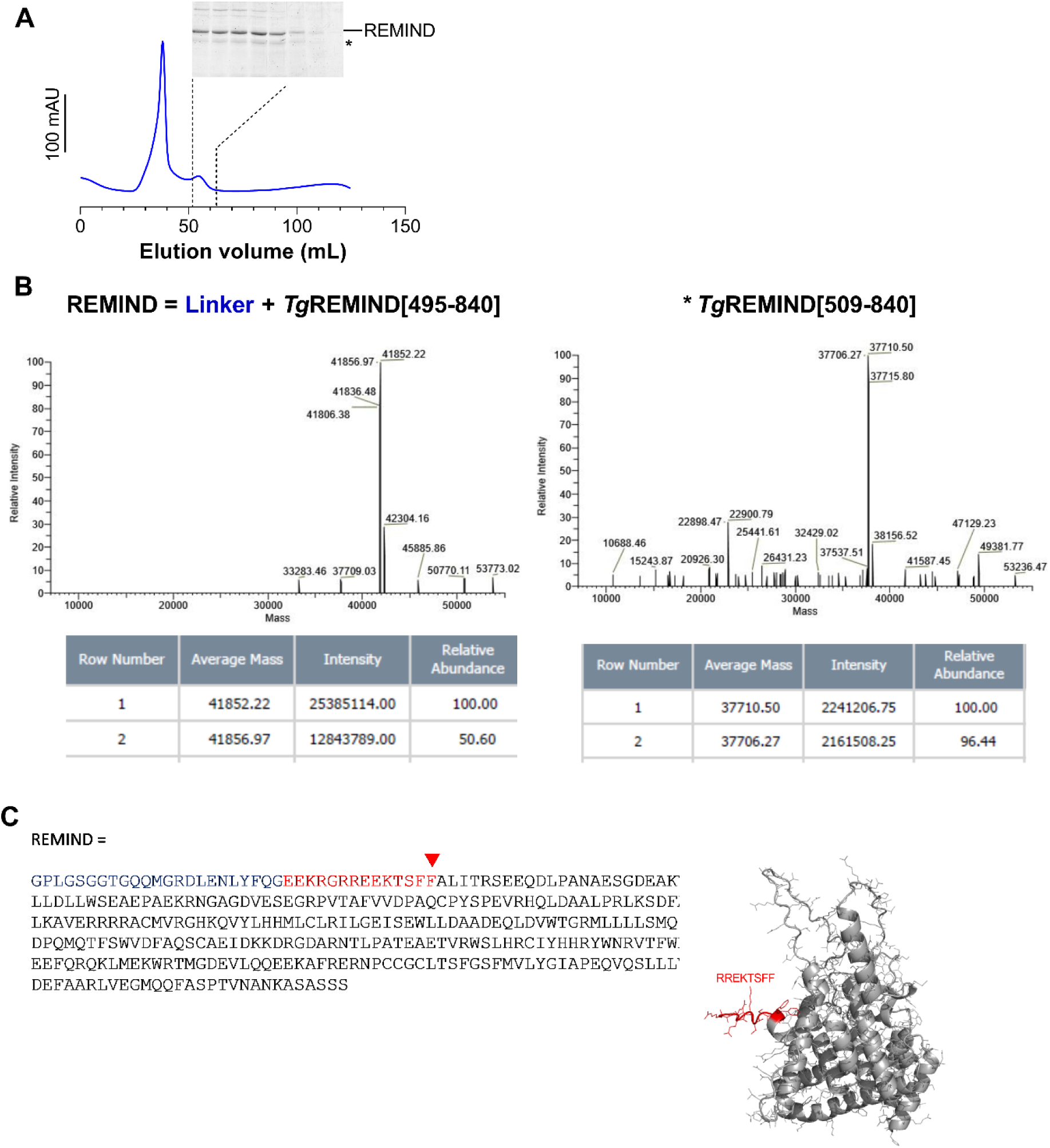
Purification of the REMIND construct. **(A)** Elution profile of REMIND during its purification by SEC with an SDS-PAGE analysis of the fractions that contain the protein of interest. The star denotes the presence of a contaminant that likely corresponds to a shorter form of the REMIND construct. AU: arbitrary unit **(C)** Mass spectrometry analysis of the REMIND batch showing the presence of a major species corresponding to REMIND (i.e., *Tg*REMIND[495-840] region fused to an N-terminal linker region] and a second one that likely corresponds to a shorter form of the construct (*Tg*REMIND[509-840) deriving from a cleavage between residue 508 and 509, as observed in (B). **(D)** Analysis of the structural model of REMIND established by AlphaFold suggests that the residues downstream of position 509 are not part of the core structure of REMIND structure and, thus, that the REMIND construct and its shorter form should have the same features.

**Supplementary Figure 4.**
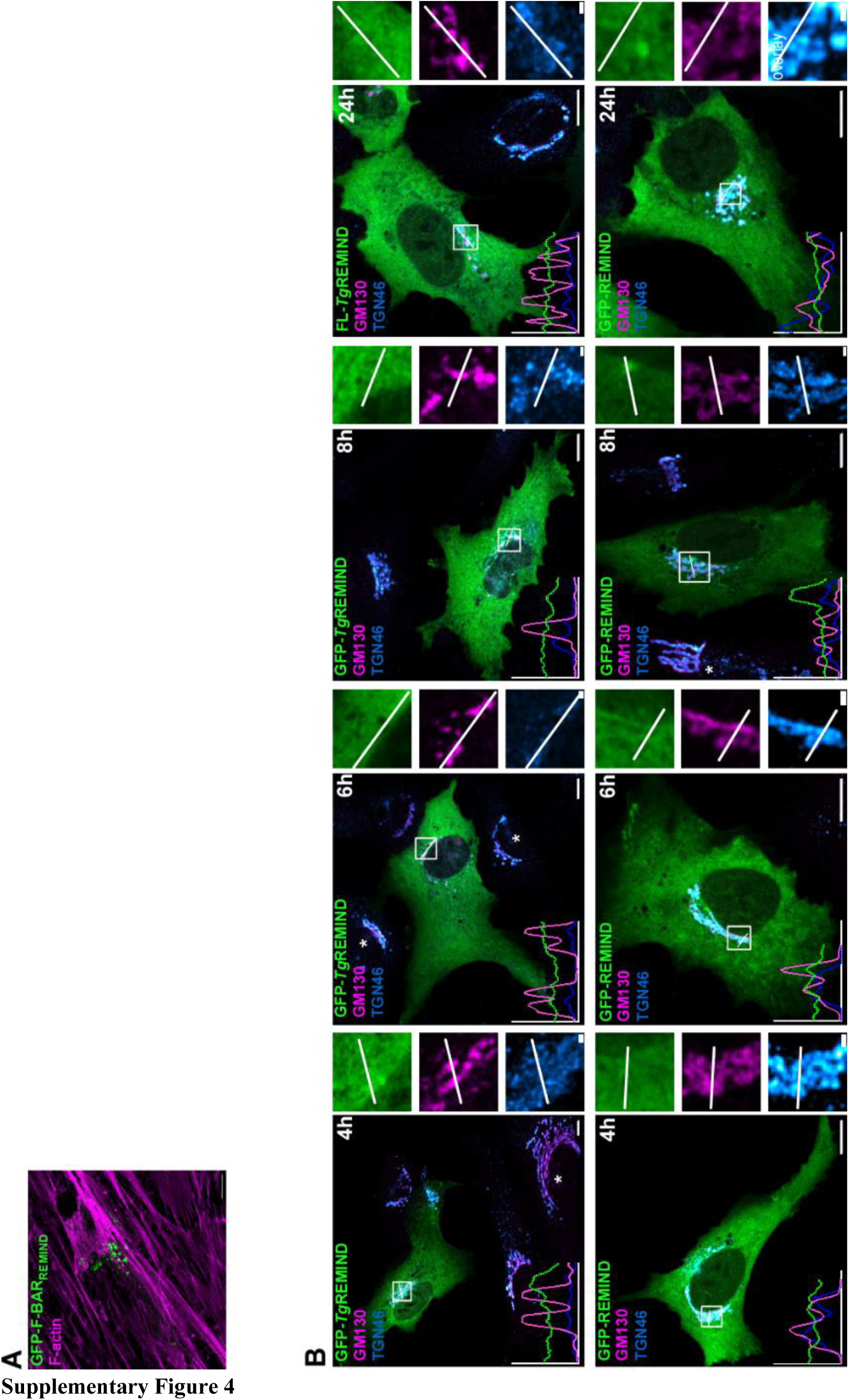
Addition data on the expression of F-BAR_REMIND_, *Tg*REMIND, and REMIND in cells. **(A)**. F-BAR_REMIND_ was expressed for 24 h in RPE-1 cells. Before observation by confocal microscopy (Leica TCS SP8, 63× NA 1.4), the cells were stained with phalloidin (ThermoFisher). Scale bar = 10 µm. **(B)** Localization of GFP-*Tg*REMIND and GFP-REMIND in RPE- 1 cells at different time points (4, 6, 8, or 24 h) after cell transfection. Before observations by confocal microscopy, cells were fixed and then labeled with an anti-GM130 antibody (magenta) and anti-TGN46 antibody (blue). Linescan shows fluorescence intensities of the green and magenta or blue channels along the white arrows shown in the insets. Stars indicate the presence of a standard Golgi apparatus in non-transfected cells. Scale bars = 10 µm (inset, 1 µm).

**Supplementary Figure 5.**
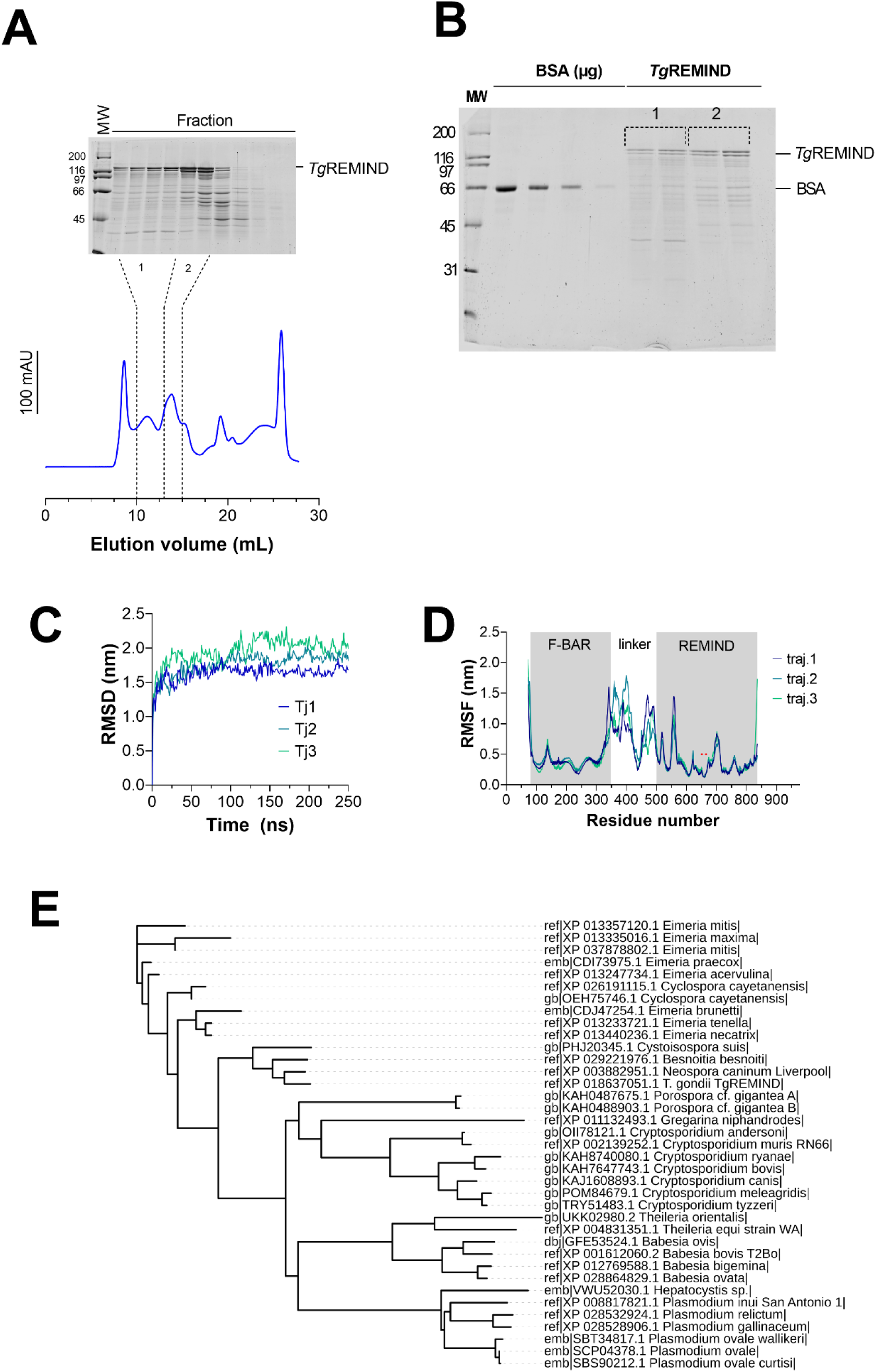
Characterization of *Tg*REMIND. (A) Purification of *Tg*REMIND. Elution profile of *Tg*REMIND during its purification by SEC with an SDS-PAGE analysis of the fractions that contain the protein of interest AU: arbitrary unit. (B) SDS-PAGE analysis showing the purity of the protein batch separated into two pools (0.3 µM and 0.9 µM stock solution for pools 1 and 2, respectively) (C) RMSD of the Cα atoms with respect to the starting and equilibrated structure of the *Tg*REMIND domain as a function of time for three 250-ns long MD trajectories. (D) RMSF values of atomic positions of Cα atoms are shown as a function of residue number. The boundaries of the F-BAR and REMIND domains and the linker between these two domains are indicated. (E) Phylogenetic analysis of *Tg*REMIND protein sequence across various Apicomplexan species.

**Supplementary Figure 6.**
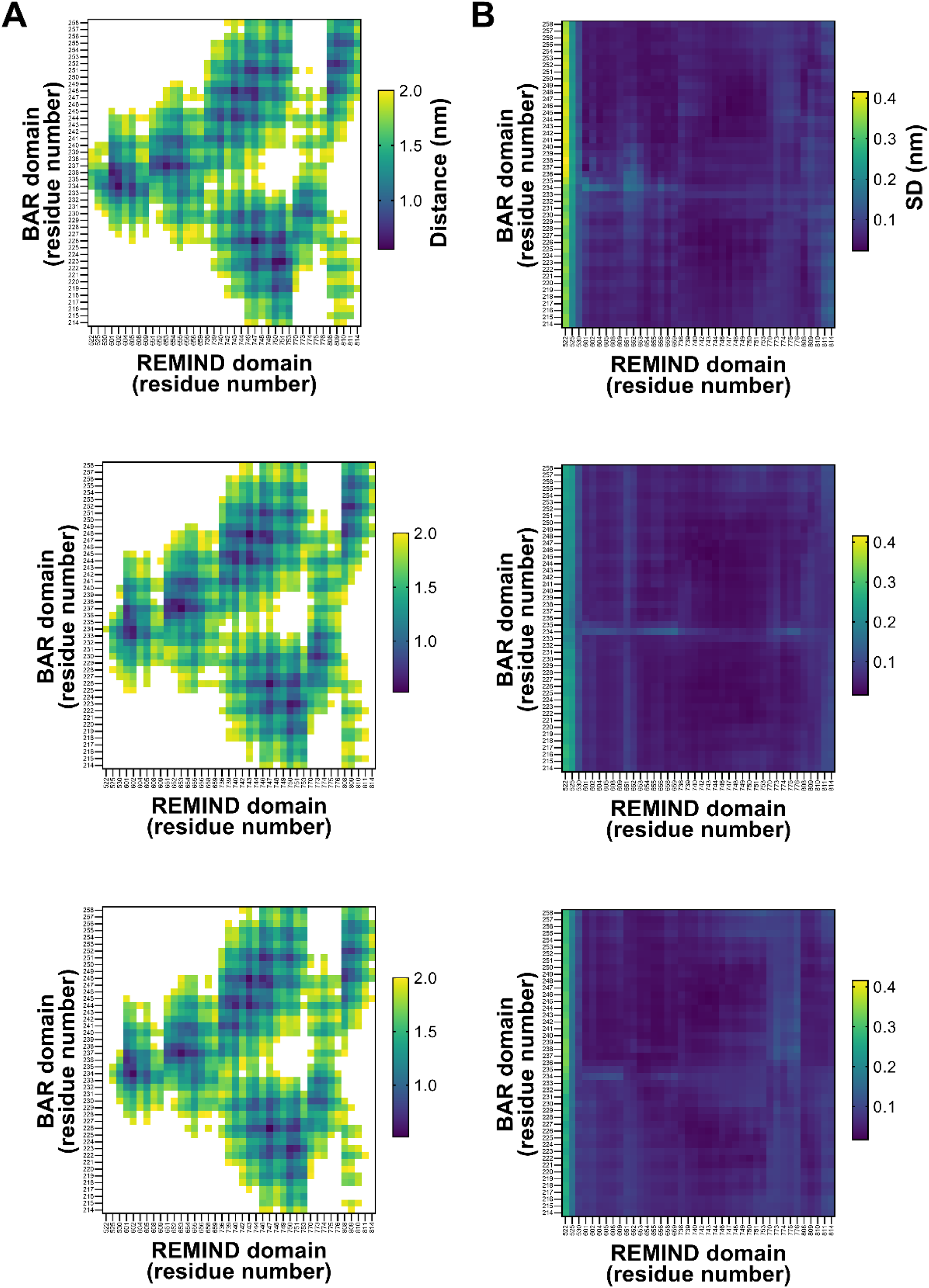
Stable association between the F-BAR and REMIND domains. **(A)** Heat maps based on a proximity matrix show that the 214-258 region of *Tg*REMIND (the extremity of the F-BAR domain) is near different residues of the REMIND domain during three independent MD trajectories. Values higher than 2 nanometers are not shown. **(B)** A second heat map, based on the standard deviation of the distance calculated for each pair of residues, shows that the BAR and REMIND domains remain stably associated.

**Supplementary Figure 7.**
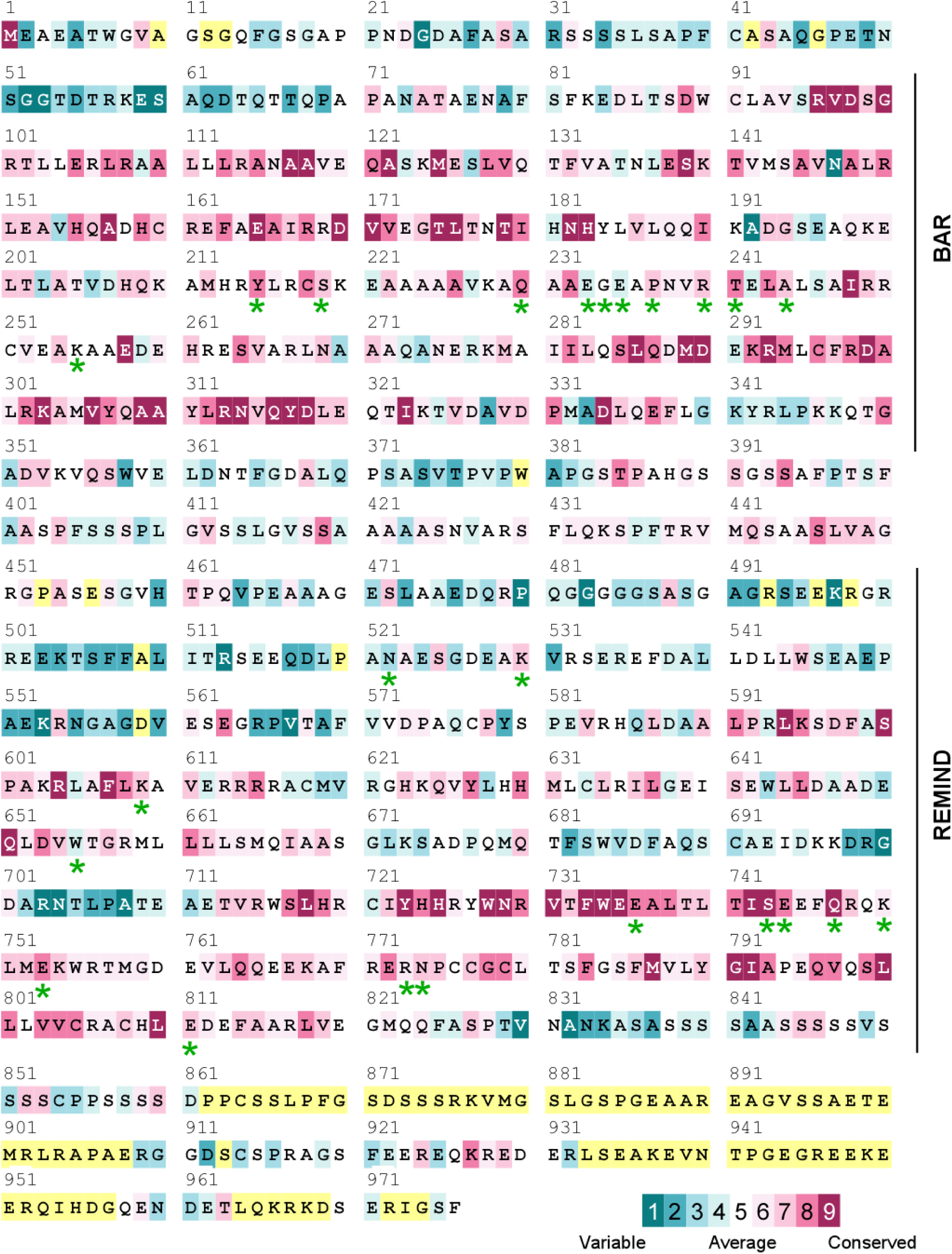
*Tg*REMIND sequence with conservation score. The conservation scale is shown. Residues for which there is insufficient data to estimate a conservation score are colored in yellow. A green star indicates the Residues responsible for forming hydrogen bonds between the BAR and REMIND domains.

**Supplementary Figure 8.**
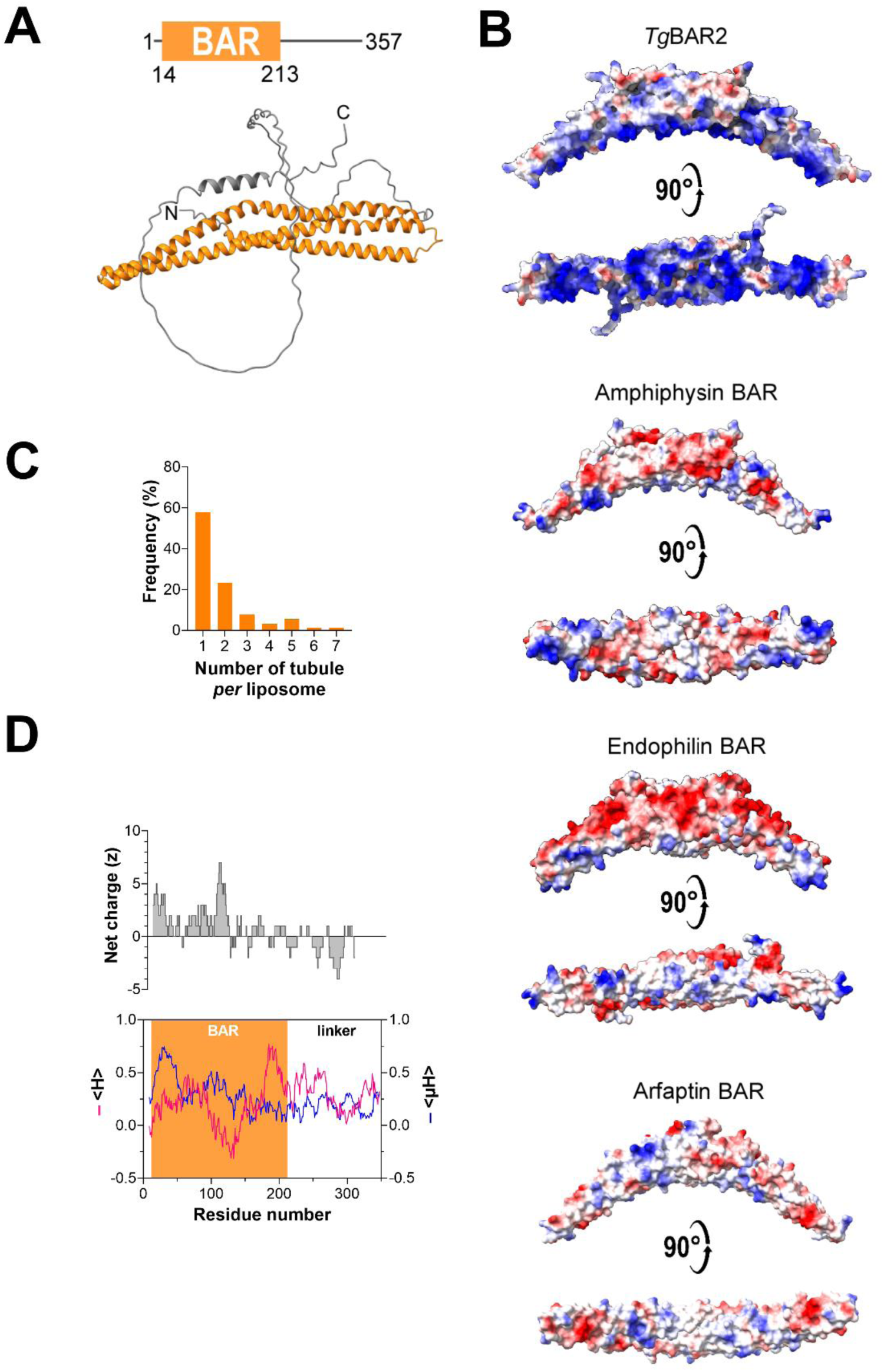
Characterization of *Tg*BAR2. **(A)** Structural organization and three-dimensional model of *Tg*BAR2 represented in ribbon mode with the BAR domain and the linker region colored in orange and grey, respectively. The position of the N- and C-terminal ends is shown. **(B)** Electrostatic surface of the BAR domain of *Drosophila* amphiphysin (1URU), human endophilin (1X03), and human arfaptin (1I49) proteins, known to impose curvature to anionic membranes. **(C)** Number of tubules *per* liposomes (considering only deformed liposomes, n = 60). **(D)** Analysis of the *Tg*BAR2 sequence shows that the linker region does not contain any 18-aa segment combining a high mean hydrophobicity (<H>), high hydrophobic moment (<µH>), and net positive charge (z) able to fold into a membrane-binding amphipathic helix. The boundaries of the BAR domain are indicated.

